# Drivers of genomic landscapes of differentiation across *Populus* divergence gradient

**DOI:** 10.1101/2021.08.26.457771

**Authors:** Huiying Shang, Martha Rendón-Anaya, Ovidiu Paun, David L Field, Jaqueline Hess, Claus Vogl, Jianquan Liu, Pär K. Ingvarsson, Christian Lexer, Thibault Leroy

## Abstract

Speciation, the continuous process by which new species form, is often investigated by looking at the variation of nucleotide diversity and differentiation across the genome (hereafter genomic landscapes). A key challenge lies in how to determine the main evolutionary forces at play shaping these patterns. One promising strategy, albeit little used to date, is to comparatively investigate these genomic landscapes as a progression through time by using a series of species pairs along a divergence gradient. Here, we resequenced 201 whole-genomes from eight closely related *Populus* species, with pairs of species at different stages along the divergence gradient to learn more about speciation processes. Using population structure and ancestry analyses, we document extensive introgression between some species pairs, especially those with parapatric distributions. We further investigate genomic landscapes, focusing on within-species (nucleotide diversity and recombination rate) and among-species (relative and absolute divergence) summary statistics of diversity and divergence. We observe highly conserved patterns of genomic divergence across species pairs. Independent of the stage across the divergence gradient, we find support for signatures of linked selection (i.e., the interaction between natural selection and genetic linkage) in shaping these genomic landscapes, along with gene flow and standing genetic variation. We highlight the importance of investigating genomic patterns on multiple species across a divergence gradient and discuss prospects to better understand the evolutionary forces shaping the genomic landscapes of diversity and differentiation.

## Introduction

Understanding the evolutionary forces that drive the accumulation of genetic diversity within and among species remains a central goal of evolutionary biology. Advances in genomic sequencing have led to numerous population genomic studies documenting genetic variation in genome scans across entire genomes (hereafter genomic landscapes). Most frequently, these studies point to a highly heterogeneous nature of these landscapes, leading to further investigations into the evolutionary forces responsible for genomic regions of elevated and reduced differentiation between diverging populations or species (Ellegren, et al. 2012; Martin, et al. 2013; Lamichhaney, et al. 2015; Vijay, et al. 2016; Sendell-Price, et al. 2020). Although theory explains how mutation, drift, selection and gene flow may shape individual features in genomic landscapes; key challenges lie in understanding the relative importance of these forces and the ‘evolution’ of genomic landscapes as species and their genomes continue to diverge over time.

Many genome scans identify regions of elevated genetic differentiation relative to the genomic background, often referred to as ‘genomic islands’, ‘differentiation islands’ or ‘speciation islands’. Theory and empirical studies support the idea that these islands can form around loci underlying local adaptation and/or reproductive isolation (e.g. Tavares, et al. 2018; McCulloch, et al. 2020; Hu, et al. 2022). Thus, delineating differentiation islands has recently become a major topic of research in the field of speciation and adaptation genomics (Burri 2017b; Martin and Jiggins 2017; Ravinet, et al. 2018; Tavares, et al. 2018; Stankowski, et al. 2019). Such investigations are best suited for groups still experiencing interspecific gene flow, *i.e*., species diverging under an isolation-with-migration or a secondary contact scenario (Harrison and Larson 2016; Roux, et al. 2016; Wolf and Ellegren 2017; Leroy, et al. 2020; Yamasaki, et al. 2020). Genomic regions containing barrier loci are more resistant to gene flow and are therefore expected to show higher levels of differentiation (the islands) as compared to the rest of the genome (the sea level, Wu 2001). For example, a number of empirical studies have demonstrated the joint role of gene flow and selection in shaping variation of the differentiation levels across the genome and identified reproductive isolation genes in regions of high differentiation (*e.g*. Tavares, et al. 2018; Martin, et al. 2019; Stankowski, et al. 2019). This highlights how the forces driving differentiation islands in specific genomic regions can be understood, however general patterns across entire chromosomes and how they evolve are more challenging to disentangle.

Part of the challenge lies in that genomic islands can also emerge from evolutionary forces not causally linked to reproductive barriers and speciation (Booker and Keightley 2018). Linked selection, the interaction between natural selection and genetic linkage may contribute to these diversity and differentiation landscapes. Two forms of linked selection are generally recognised: background selection and genetic hitchhiking; although their relative importance is still debated (Stephan 2010). Background selection (Charlesworth, et al. 1993), the effect of natural selection against deleterious alleles at linked neutral polymorphism, is known to reduce diversity, particularly in regions with relatively high gene density (Corbett-Detig, et al. 2015; Wolf and Ellegren 2017). Similarly, due to genetic hitchhiking, neutral alleles are dragged along with positively selected ones (Smith and Haigh 1974). Linked selection reduces effective population size (*N_e_*) and can lead to regions of decreased diversity and elevated relative differentiation. In regions of low recombination, linked selection can generate footprints that extend over larger genomic regions around the positively or negatively selected loci (Charlesworth and Campos, 2014). Thus, nucleotide diversity (*e.g., π*) and relative differentiation (*e.g., F_ST_*) estimates are expected to be negatively correlated (Burri 2017a). Such correlations have been reported in *Ficedula* flycatchers (Burri, et al. 2015), *Heliconius* (Edelman, et al. 2019; Martin, et al. 2019; Van Belleghem, et al. 2021), *Helianthus* sunflowers (Renaut, et al. 2013), the Pacific cupped oyster (Gagnaire, et al. 2018), warblers (Irwin, et al. 2018), and hummingbirds (Henderson and Brelsford 2020). These findings suggest that a negative correlation between diversity and relative differentiation taken in context to the local recombination rate across chromosomes, provides a useful null expectation for interpreting genomic landscapes of divergence.

Given the range of causes of heterogeneity in genomic landscapes, that may also depend on divergence time between species compared, an important goal of speciation research is therefore understanding how genomic landscapes ‘evolve’ and accumulate differences over time. However, a major limitation of most genomic landscapes lies in the limited number of pairwise comparisons available. Whether genomic divergence evolves in a predictable way and patterns of divergence are conserved among different species pairs, is also difficult to determine when comparing individual studies from different genera and families. Comparing independent genome scan studies may potentially show different snapshots along the divergence gradient relevant to their respective genera/family, confounded with contrasting genomic features such as recombination rate, effective population size and demographic history. One approach to address this problem is to investigate these genomic landscapes along a progression through time by using a series of species pairs along a divergence gradient within a genus. With multiple pairwise comparisons along a divergence gradient, this allows for tests of correlations in genomic measures expected under different evolutionary scenarios (see Box 1) whilst controlling for similar genomic backgrounds. To understand the processes behind the heterogeneous differentiation landscapes along a divergence gradient, a suite of summary statistics, widely used in population genomics, can be employed (Han, et al. 2017; Irwin, et al. 2018). By comparing local genomic patterns and genomic wide correlations between summary statistics along the divergence gradient (Box 1), we can begin to identify the tempo of differentiation and ask whether differentiation is accumulated in a predictable way in independent lineages and if these patterns are conserved along regions of the genome.

### Box 1: Correlations of genomic landscapes under different scenarios of divergence

Following Han *et al* (2017) and Irwin *et al* (2018), four main evolutionary scenarios can be hypothesized. The first scenario is ‘divergence with gene flow’ where selection at loci contributing to reproductive isolation restricts gene exchange between diverging species, locally elevating genomic differentiation (leading to both high *F_ST_* and *D_XY_*) and reducing genetic diversity. The second scenario is ‘allopatric selection’ where linked selection occurs independently within each species after the split leading locally to lower *π* and higher *F_ST_*.Allopatric selection has opposite effects on *D_XY_*, leaving it relatively unchanged. The third scenario is ‘recurrent selection’ where the same selective pressure reduces diversity at selected and linked loci leading to lower polymorphism within populations but similar divergence, *i.e*.relatively low *π* and *D_XY_* due to its dependence on ancestral polymorphism and high *F_ST_*. The fourth and last scenario is ‘balancing selection’ where ancestral polymorphism is maintained between nascent species, resulting in elevated genetic diversity and low genetic differentiation. Then *π* is expected to be higher than neutral (as is *D_XY_*, due to the high ancestral diversity) while *F_ST_* is expected to be low.

**Box 1 Figure:**
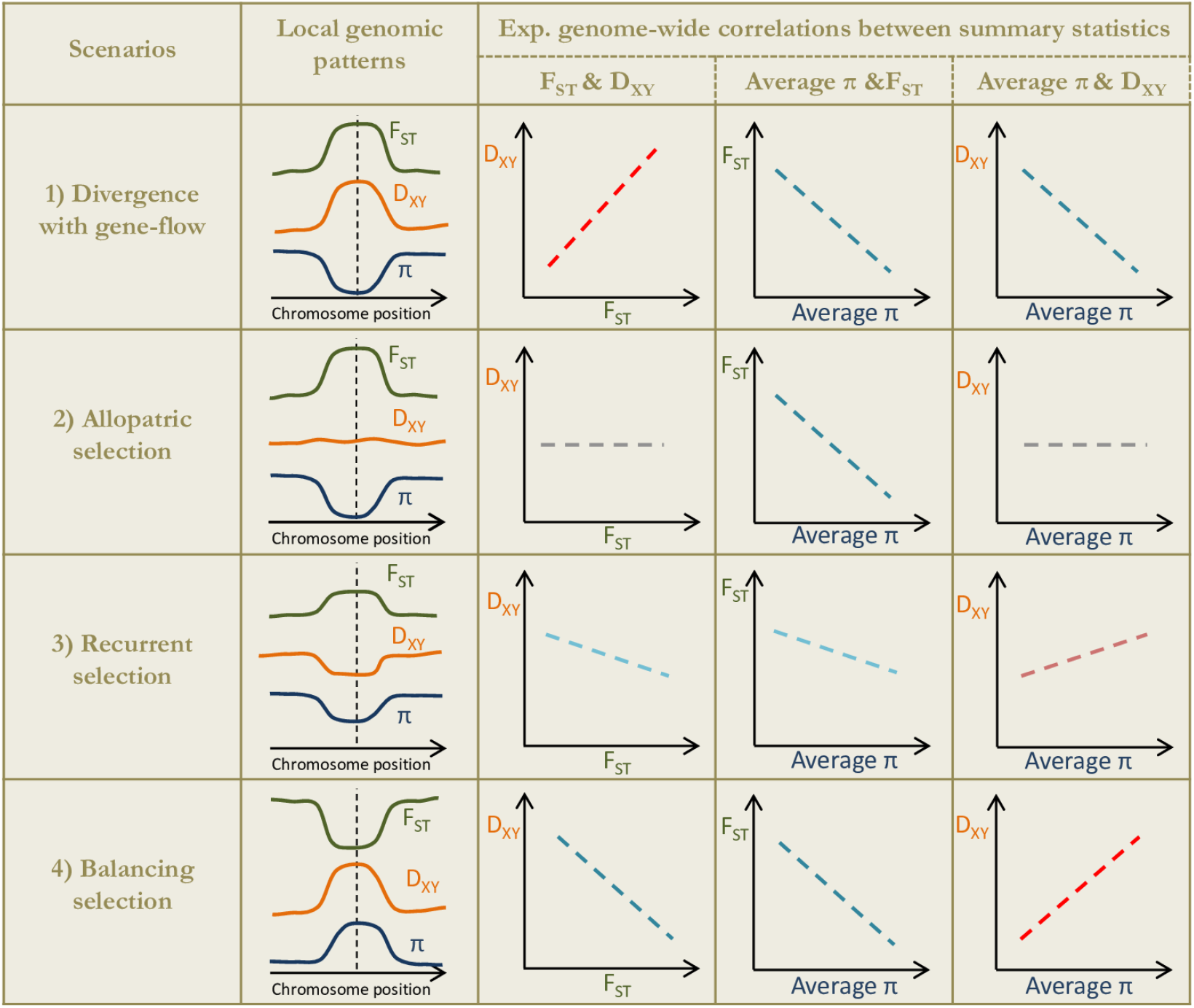
Expected correlations of pairs of summary statistics describing the genomic landscapes of diversity and differentiation associated with the four different scenarios proposed by Han et al (2017) and Irwin et al (2018). Positive and negative relationships are shown in red and blue, respectively. In the second column, local patterns associated with each scenario are described, i.e. (1) divergence with gene flow (a reproductive barrier to gene flow), (2) allopatric selection (a selective sweep in one of the two populations), (3) recurrent selection (a footprint of ancestral and still ongoing selection), (4) balancing selection. Average π corresponds to the averaged value of π for the two species included in the pairwise comparison. Given that both π and population-scaled recombination rates (ρ) are dependent on N_e_, similar relationships are expected for the relation with F_ST_ or D_XY_ and ρ, as compared with π. It is important to note that the contrast is also dependent on the divergence time (not shown here), with more distinct F_ST_ peaks above the genomic background in the least divergent pairs. Note that scenarios three and four are associated with the same directions of the correlations but are expected to have very different local signatures for F_ST_, D_XY_ and ρ, therefore allowing the distinction of these last two scenarios.

In this study, we focused on white poplars and aspens from the section *Populus* within the genus *Populus*. These trees are widely distributed in Eurasia and North America (Supporting Fig. S1 and Table S1) and provide a set of species pairs along the continuum of divergence. Divergence times among species pairs vary from 1.3 to 4.8 million years ago (Shang, et al. 2020). This provides an excellent system to investigate the evolution of genomic landscapes of diversity and divergence through time and to better understand the relative contribution of different evolutionary processes to genomic landscapes. We use whole-genome resequencing data from eight *Populus* species (Supporting Fig. S1 and Table S1) to address the following questions: (1) How do genomic landscapes of differentiation accumulate along the divergence gradient? (2) Are differentiation patterns across the genomic landscape repeatable among independent lineages? (3) What are the main evolutionary processes driving these heterogeneous landscapes of diversity and differentiation along the divergence gradient? (4) Which divergence scenario is consistent with ‘differentiation islands’ in each species pair?

## Results and Discussions

### Strong interspecific structure despite interspecific introgression

A large dataset of 30,539,136 high-quality SNPs was obtained by identifying SNPs among individuals from seven *Populus* species (after the exclusion of *P. qiongdaoensis*, see Materials and Methods). Neighbor-joining (Fig. 1a) and admixture analyses (based on a subset of 85,204 unlinked SNPs, Fig. 1b and Supporting Figs. S2 and S3) identified seven genetic groups, which were consistent with previously identified species boundaries based on phylogenomic analyses (Shang et al, 2020). Additionally, Admixture also indicated potential introgression between the subtropical species *P. adenopoda* and two recently diverged species, *P. davidiana* and *P. rotundifolia* (Fig. 1b and Supporting Fig. S3).

**Figure 1.**
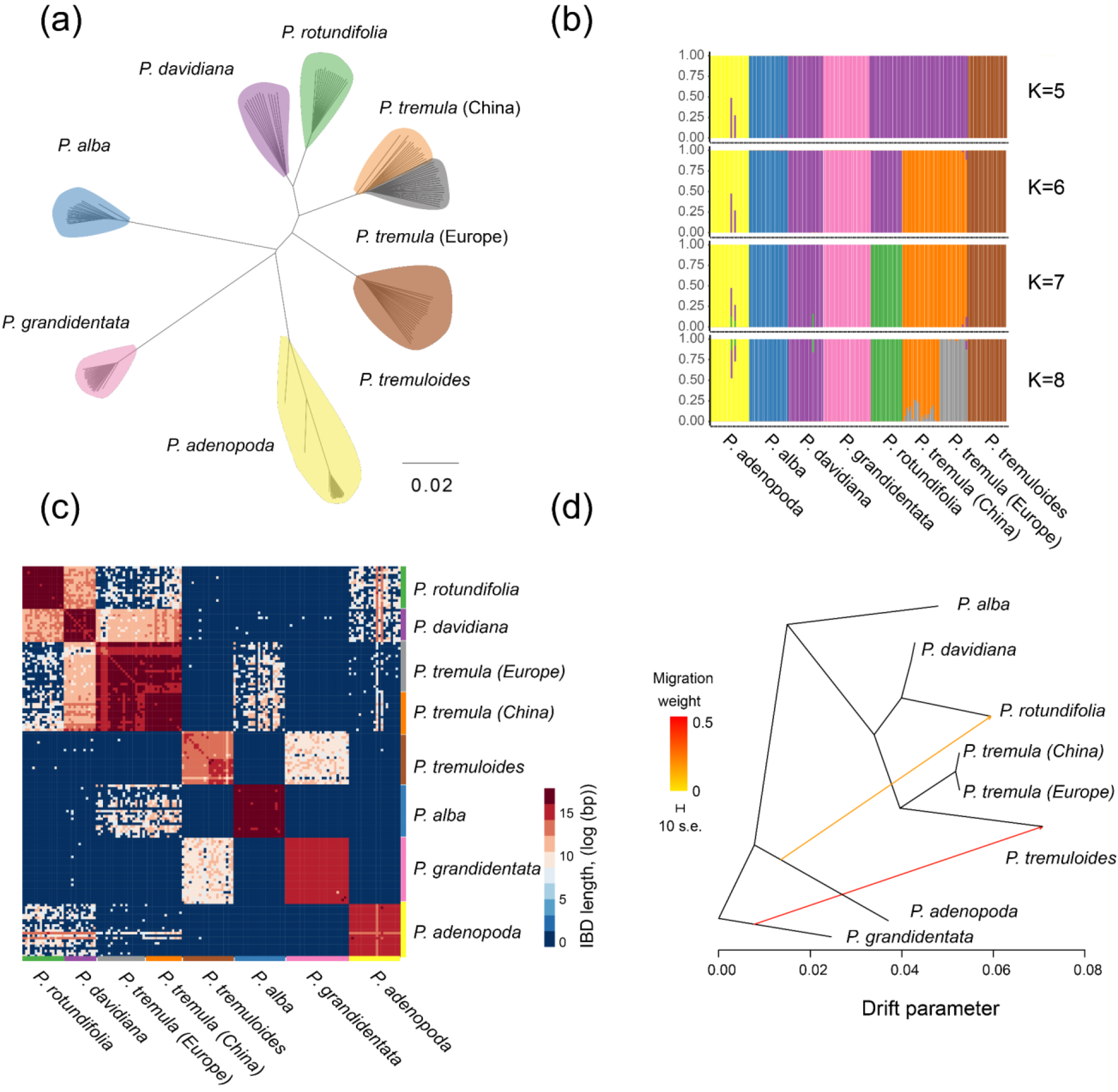
Genetic structure and evolutionary relationships among Populus (poplar and aspen) accessions. **(a)** Neighbor-joining tree based on all SNPs for seven Populus species. Colored clusters represent different species according to legend. **(b)** Estimated membership of each individual’s genome for K = 5 to K = 8 as estimated by Admixture (best K = 7). **(c)** Identity by descent (IBD) analysis for seven Populus species. Heatmap colours represent the shared haplotype length between species. **(d)** The maximum likelihood tree inferred by TreeMix under a strictly bifurcating model with two migration events.

Identity-by-descent (IBD) analyses (Fig. 1c) also identified seven reliable clusters, corresponding to the same species boundaries, but further pinpointed some shared haplotypes among the aspen species *P. davidiana*, *P. rotundifolia* and *P. tremula*, indicating recent introgression or incomplete lineage sorting among these species. The IBD results also provide support for extensive introgression between two pairs of highly divergent species with overlapping distributions, including *P. alba* and *P. tremula*, and also *P. grandidentata* and *P. tremuloides*. These results suggest a scenario of divergence with past and/or ongoing gene flow for some species pairs, either due to isolation-with-migration or secondary contact. Given that hybrids between these species can be obtained in nature and/or obtained through artificial breeding (Vanden Broeck, et al. 2005; Deacon, et al. 2017; Deacon, et al. 2019), this suggests that reproductive isolation remains incomplete even after substantial divergence times (net divergence *d_a_*: 0.023 for *P. alba* - *P. tremula; d_a_*: 0.025 for *P. tremuloides* - *P. grandidentata*).

We have further confirmed that the tree topology recovered with TreeMix (Pickrell and Pritchard 2012) was consistent with phylogenetic relationships found in a previous study (Shang, et al. 2020). This expected main topology explained 95.8% of the total variance under a drift-only model of divergence. In addition, TreeMix was used to infer putative migration events in *Populus* (Fig. 1d). Adding a single migration edge allowed us to account for 98.9% of the total variance (Supporting Fig. S4). This event was inferred from *P. grandidentata* to *P. tremuloides* and is consistent with previous reports of extensive hybridization and introgression between these two species (Dickmann 2001; Deacon, et al. 2019). A second migration edge was inferred from *P. adenopoda* to *P. rotundifolia*, which allowed us to explain 99.6% of the total variance (Fig. 1d). By adding more migration edges, the variance explained plateaued (increasing by less than 0.1%, which was considered as too marginal, Supporting Fig. S4). Therefore, we considered the bifurcating tree with two migration events as the best scenario in this analysis explaining the historical relationships among these *Populus* species based on our data and sampling.

### Detecting local genomic patterns consistent with the four scenarios

Using non-overlapping 10kb sliding windows spanning the genome, we reported diversity and divergence estimates for all species and species pairs (Fig. 2a). Mean *F_ST_* varied from 0.23 between *P. davidiana* - *P. rotundifolia* to 0.71 between *P. adenopoda* - *P. grandidentata*, whereas *D_XY_* (Supporting Fig. S5) ranged from 0.016 (*P. davidiana* - *P. rotundifolia*) to 0.028 (*P. adenopoda* - *P. grandidentata*). Looking at the genomic landscapes of *F_ST_*, the observed variation correlates with the levels of introgression for species pairs exhibiting with expected hybridization and/or previously detected migration events under TreeMix (Supplementary Note 1, Supporting Fig. S6). The average *π* varied from 3.5×10^-3^ in *P. grandidentata* to 8.4×10^-3^ in *P. tremuloides* (Fig. 2a). The relatively high average *π* value observed in *Populus* species is consistent with the large SMC++-inferred effective population sizes for these species (Supplementary Note 2, Supporting Fig. S7 and S8) and the fast LD decay (Supporting Fig. S9). In addition to nucleotide diversity and differentiation across 10kb sliding windows, we also computed *ρ* for each species. Although the level of genetic diversity was low in *P. grandidentata*, the *ρ* of this species was the highest. Correlation analysis between π and ρ showed that all species except *P. grandidentata* showed significantly positive correlations (Supporting Fig. S10), while we detected significant negative correlations between gene density and *π* in all *Populus* species (Supporting Fig. S11).

**Figure 2.**
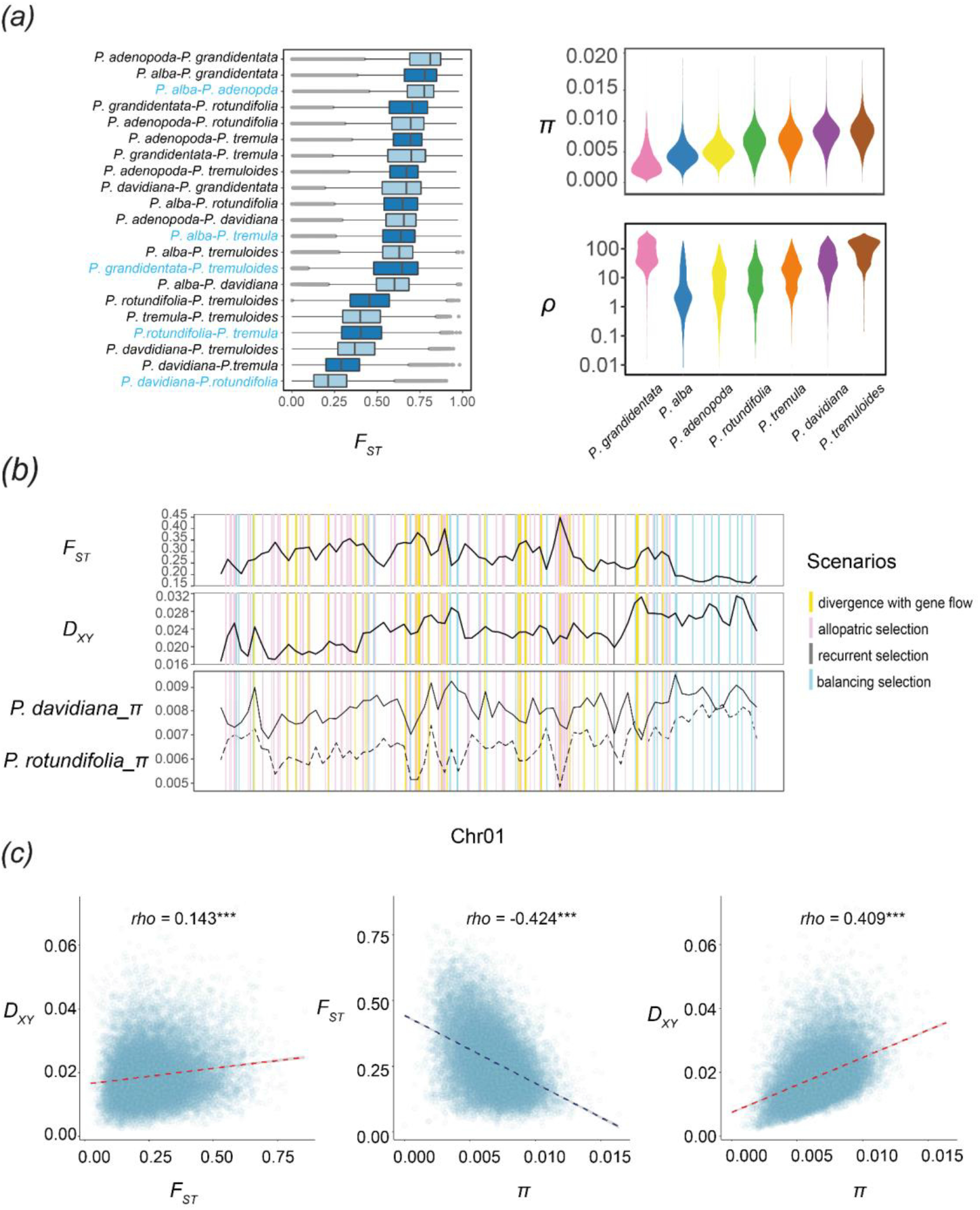
**(a)** Observed variance in F_ST_ for all species pairs, and π and ρ for seven Populus species, calculated across 10kb windows. The five representative species pairs were labeled in blue. Note that the unit of ρ is 4Ner and that ρ is log-scaled. **(b)** Landscapes of π, F_ST_, and D_XY_ on chromosome 1 for the two closest species, P. davidiana and P. rotundifolia. **(c)** Genome-wide correlation analysis for π, F_ST_, and D_XY_ between P. davidiana and P. rotundifolia. P values less than 0.001 are summarized with three asterisks.

We then identified regions that could be consistent with the alternative divergence scenarios (described in Box 1) for five representative species pairs (Supporting Fig. S12). These species pairs were selected to represent distinct stages across the divergence gradient, from early to late stages of speciation (blue labels in Fig. 2a). Our results are consistent with a heterogeneous distribution of the four scenarios along the genome for all five species pairs (Fig. 2b, see also Supporting Fig. S13-S17 and Table S2). This generates different genome-wide patterns of correlations among species pairs, rather than a single scenario at play across the whole genome (Fig. 2c, Box 1). The majority of the detected genomic windows (74.3%-78.7%) in all five species pairs fit a scenario of “allopatric selection”, in which the excess of *F_ST_* was driven by low *π* and not higher *D_XY_* (pink bars in Fig. 2b, Supporting Fig. S13-S17 and Table S2). Such a signature is consistent with recent footprints of positive or background selection on genomic differentiation and is therefore consistent with the hypothesis of a prime role of linked selection (see also Supplementary Note 3 for an explicit detection of selective sweeps). Genomic regions fitting the scenario of ‘balancing selection’ (scenario d in Supporting Fig. S12) are the second most frequent for all investigated species pairs (11.6%-13.9% of detected regions). This scenario is characterized by an elevated *D_XY_* but a low *F_ST_* implying the action of balancing selection in shaping the heterogeneous landscape of divergence. In addition, we found support for divergence-with-gene flow in all five species pairs (5.5%-8.1%), suggesting that genomic heterogeneity in the levels of gene flow due to species barriers play a role in shaping genomic differentiation landscapes. Interestingly, this result holds true for all five species pairs we investigated in detail, *i.e*., regardless of the level of gene flow or the stage along the *Populus* speciation continuum. Indeed, limited gene flow was inferred between *P. adenopoda* - *P. alba*, but regions with high *D_XY_* were also identified in this highly diverged species pair and could be rather due to shared ancestral polymorphisms. In contrast, at the early stage of divergence, local barriers to gene flow may play an important role in genomic heterogeneous divergence, as significantly positive correlations between *D_XY_* and *F_ST_* are found (Fig. 2c), which is consistent with a divergence with gene-flow scenario (Box 1). Besides, for early stages of divergence background selection may have too limited power to explain alone regional patterns of accentuated differentiation (Burri 2017b).

### Genomic landscapes of variation across the continuum of divergence

We calculated genome-wide correlations of divergence, nucleotide diversity, and recombination across non-overlapping 10kb windows spanning the whole genome between pairwise comparisons of species or species pairs. The degree of correlation of both the relative and absolute divergence landscapes between pairs of species supports a highly conserved pattern among the five investigated species pairs (Fig. 3a-b, between *P. adenopoda -P. alba* and *P. tremula -P. alba* for *F_ST_* and between *P. rotundifolia - P. davidiana* and *P. rotundifolia - P. tremula* for *D_XY_*). The correlations of *F_ST_* landscapes become stronger when the overall differentiation increases along the divergence gradient (Fig. 3a), whereas correlations of *D_XY_* remained similar (Fig. 3b). For instance, the correlation of *F_ST_* between *P. tremula* - *P. alba* and *P. rotundifolia - P. davidiana* is 0.24, while the value for the two most divergent species pairs (between *P. adenopoda* - *P. alba* and *P. tremuloides* - *P. grandidentata*) is 0.56. This suggests that the effect of linked selection accumulates as differentiation accumulates. Comparing landscapes of the nucleotide diversity *π* between species (Fig. 3c), we observed that the correlation coefficients vary substantially, from 0.16 (*P. tremula* versus *P. grandidentata*) to 0.52 (*P. rotundifolia* versus *P. davidiana*). The correlation generally decreases with the phylogenetic distance. We notably reported the strongest correlation coefficient for the phylogenetically closest pair of species: *P. rotundifolia* and *P. davidiana* (Fig. 3c). Pairwise comparisons of the local recombination rates inferred independently for all species also revealed only positive correlations (Fig. 3d), with the highest positive correlation coefficient of *ρ* again observed between the two closest related species, *P. davidiana* and *P. rotundifolia* (0.47), while the weaker correlation was observed for *P. davidiana* and *P. grandidentata* (0.08). Most of the lower values (correlation coefficients < 0.2) were found when comparing *P. grandidentata* with other species, suggesting again a unique recombination landscape in this species (Fig. 2a). Interestingly, correlations of *π* were in general higher than those of *ρ*, indicating that not only recombination rate variation shapes nucleotide diversity. Overall, landscapes of genetic diversity, divergence and recombination rate remain relatively stable across different species or species pairs (Fig. 3), which implies relatively conserved genomic features across all species. This phenomenon has also been observed in few other plant and animal model systems (Nosil and Feder 2012; Renaut, et al. 2014; Burri, et al. 2015; Wang, et al. 2020).

**Figure 3.**
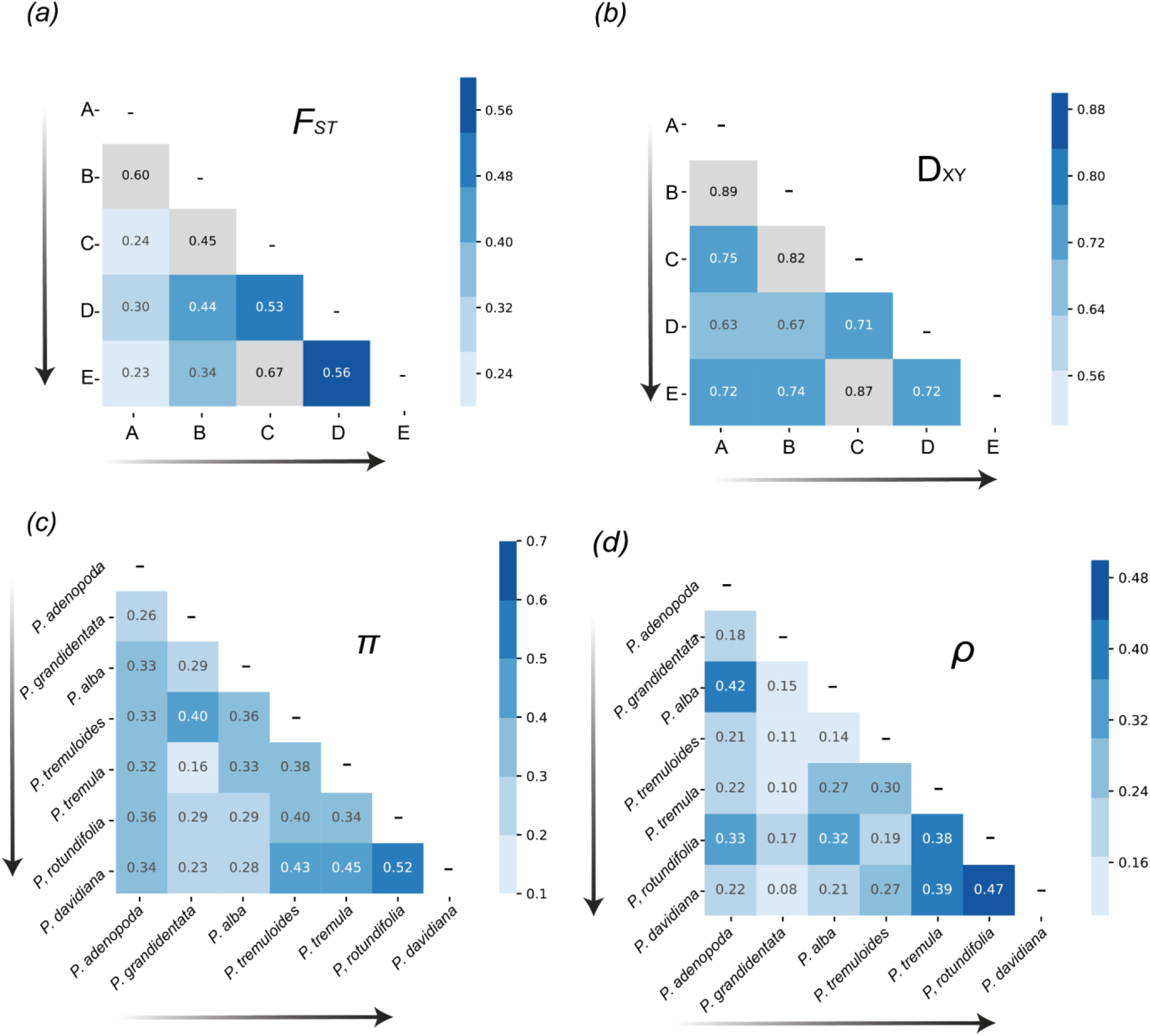
Correlations analyses of within-species diversity or among-species divergence landscapes **(a-b)** Correlation coefficients of F_ST_ or D_XY_ between species pairs. A-E are abbreviations for the five representative species pairs:A, P. rotundifolia-P. davidiana; B, P. rotundifolia-P. tremula; C, P. alba-P. tremula; D, P. grandidentata-P. tremuloides; E, P. adenopoda-P. alba. The species pairs are ordered across the divergence gradient (arrows). Comparisons containing a shared species were masked (grey squares). **(c-d)** Correlation coefficients of π or ρ between species. The order of the species is based on the order of species divergence from the root. All the values are significantly positively correlated (p < 0.001).

### Correlated patterns of genome-wide variation across the *Populus* continuum of divergence

The relatively conserved genomic patterns observed across independent species pairs is consistent with an important role for linked selection in shaping the genomic landscapes of differentiation (Fig. 3). One possibility is background selection could play a major role, as deleterious mutations are much more common than beneficial ones. Therefore, we test if background selection (BGS) may have driven these patterns alone. According to expectations proposed by Burri (2017b), the correlations between genomic variation and recombination rates are most likely to have arisen through background selection. First, the correlation between *F_ST_* and *ρ* is expected to become stronger with divergence, as lineage-specific effects of background selection accumulate with time. Second, *D_XY_* and *π* are highly correlated with one another under BGS, because diversity can be inherited from ancestors, being passed down over lineage splits. Third, *π* and *ρ* remain highly correlated, because background selection continues to play a role in the daughter populations after speciation.

The use of several species across a continuum of divergence allows us to evaluate how the correlations evolve through this continuum, from early to late stages of speciation. To this end, we used the level of genetic distance between each species pair (net nucleotide differences, *d_a_* = *D_XY_* – mean *π*) as a proxy for the divergence time and we reported linear relationships between correlation coefficients across the 21 species pairs (Fig. 4). However, we found negative relationships between *F_ST_* and *ρ*, and no significant changes associated with time since divergence (Fig. 4a). This is inconsistent with expectations under background selection (Burri 2017b). Similar investigations for *π* and *F_ST_* showed significantly negative correlations which became stronger as divergence increases (Fig. 4b). We also recovered a strong positive corre-lation between average *π* and *ρ* (Fig. 4c), and a similar trend was found between *π* and *D_XY_* (Fig. 4d). This trend is inconsistent with the general hypothesis that such correlations should remain highly correlated as divergence increases (Burri 2017b). Pairwise correlations between *D_XY_* and *F_ST_* were significantly positive across the entire divergence continuum, and these correlations tend to become weaker as divergence increases (Fig. 4e). The observed patterns differ from expectations under a simple scenario with background selection as the sole factor shaping the heterogeneous landscape of differentiation across species, indicating that additional evolutionary factors contribute to the observed signal (Burri 2017b). Our analyses indicate that extensive gene flow and incomplete lineage sorting may contribute to differentiation landscapes as well, in particular in the early stages of the speciation continuum. However, with increasing divergence, the reduced gene flow and limited shared standing genetic variation may contribute less to differentiation landscapes. Consistent with our findings, studies in monkey flowers, threespine stickleback, and in warblers also suggest that background selection may be too subtle to drive alone conserved genomic patterns across multiple species (Irwin, et al. 2018; Stankowski, et al. 2019; Rennison, et al. 2020). Indeed, these studies indicate either adaptive introgression or shared standing genetic variation also play major roles in generating similar patterns of genomic differentiation. In addition, we also detected selective sweep regions across the genome for all species, many of which were species-specific loci (Supplementary Note 3).

**Figure 4.**
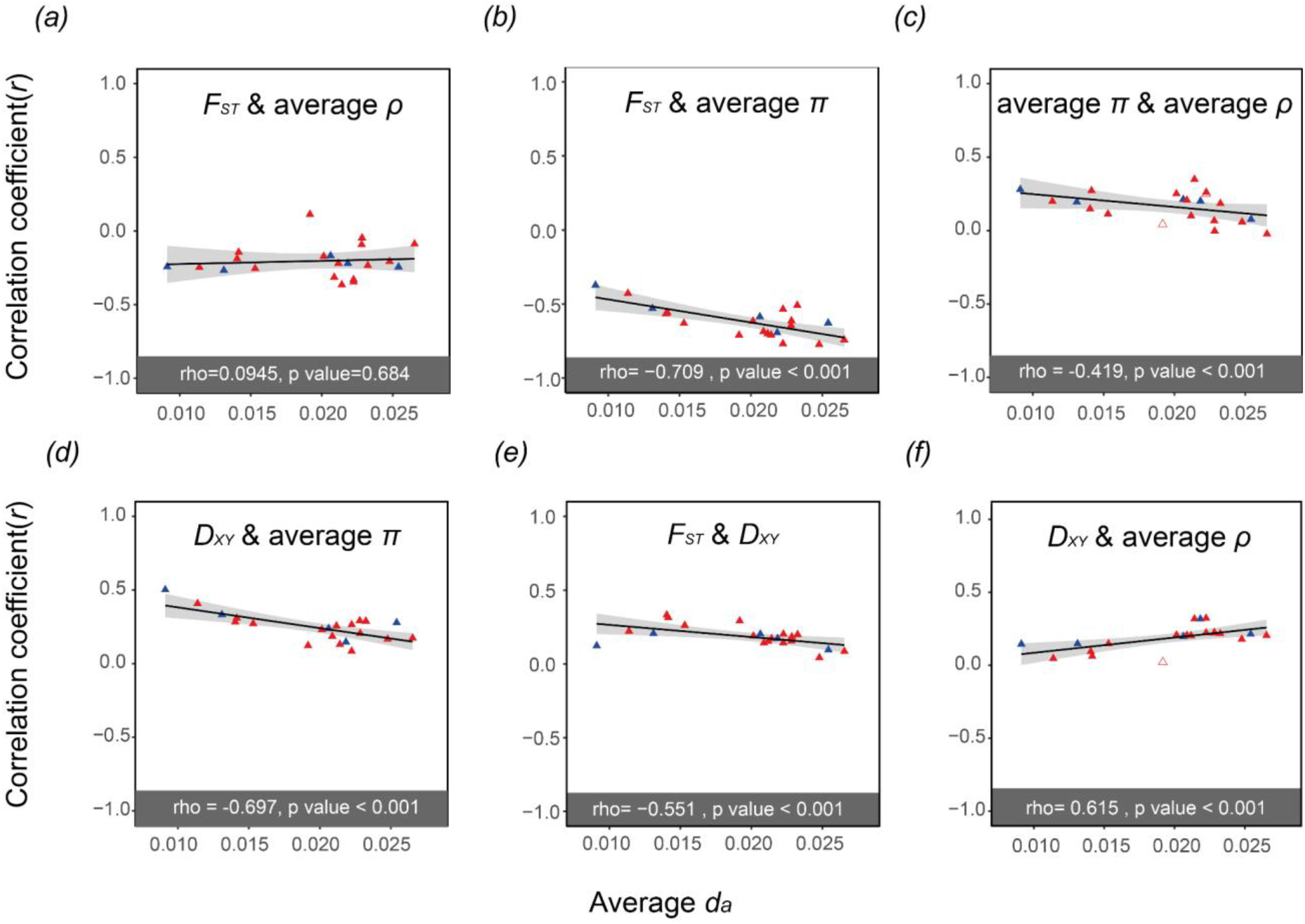
Correlations between variables for all species comparisons plotted against the averaged d_a_, used here as a measure of divergence time. The filled triangles indicate when the correlation coefficients are significant (p < 0.01). The blue triangles correspond to the correlation coefficients of the five representative species pairs shown in Fig. 2a and 3a-b. The results for all the other species pairs are shown in red. The upper panels show how the relationships between F_ST_ and **(a)** average ρ, or (**b**) average π vary for pairs of species with increasing divergence time; and between average π and average ρ for all species pairs investigated **(c)**. The lower panels show how the relationships between D_XY_ and **(d)** average π, **(e)** F_ST_, and **(f)** average ρ vary for pairs of species with increasing divergence time.

## Conclusions

In this study, we investigated the evolution of the genomic landscape across a divergence time continuum of seven species of *Populus*. By investigating evolution of diversity and differentiation landscapes across this divergence continuum, we provide a valuable case-study in terms of the number of species pairs analyzed (see also Stankowski, et al. 2019). Our analyses support the primary role of linked selection, in particular background selection in shaping the empirical patterns of genomic differentiation, but its contribution alone is not sufficient to explain the accumulation of changes in the genomic landscapes as species diverge over time. The observed positive correlations between *F_ST_* and *D_XY_* in all species pairs indicate that shared ancient polymorphism must also play a very important role. In addition, we observed extensive introgression among species with parapatric distributions, despite a high level of divergence among the most divergent hybridizing species (*d_a_* = 0.025). This is notable since the net divergence values are greater than the upper boundary for the ‘grey zone of speciation’ reported by Roux et al. (2016) for both vertebrate and invertebrate animals (*d_a_* from 0.005 to 0.02). Further investigations across divergence continua in other plant systems are needed to determine if this is a general pattern in plants, or a feature of the specific demographic and evolutionary history of the *Populus* system. In the future, the investigation of speciation along multiple species complexes, together with the inclusion of the different scenarios of selection in more sophisticated demographic modeling approaches could represent a major step forward to provide a better description of the processes at play.

## Materials and methods

### Sampling, sequencing and read processing

Species of the genus *Populus* are perennial woody plants, dioecious, and widely distributed across the Northern Hemisphere (Stettler, et al. 1996). The genus *Populus* comprises six sections containing 29 species, among which ten species form the section *Populus* (Stettler, et al. 1996; Jansson, et al. 2010). The genus *Populus* is well studied not only due to the trees’ economic and ecological importance, but also due to their small genome sizes (<500Mb), diploidy through the genus (2n = 38), wind pollination, extensive gene flow among species, and sexual and vegetative reproductive strategies (Rajora and Dancik 1992; Martinsen, et al. 2001; Suarez-Gonzalez, et al. 2016). Among all woody perennial angiosperm species, the genome of *Populus trichocarpa* was sequenced and published first (Tuskan, et al. 2006). In addition to *P. trichocarpa*, another well-annotated genome assembly is available (*P. tremula;* Schiffthaler, et al. 2019).

Two hundred and one samples were collected from eight species of *Populus* section *Populus* in Eurasia and North America (supplemental material, Fig. S1 and Table S1). The leaves were dried in silica gel first and were then used for genomic DNA extraction with Plant DNeasy Mini Kit (Qiagen, Germany). To increase the purity of total DNA, we used the NucleoSpin gDNA Clean-up kit (Macherey-Nagel, Germany). Whole genome resequencing was performed with 2 x 150bp paired-end sequencing technology on Illumina HiSeq 3000 sequencer at the Institute of Genetics, University of Bern, Switzerland.

All raw sequencing reads were mapped to the *P. tremula* 2.2 reference genome (Schiffthaler, et al. 2019) available on the PlantGenIE.org website (Sundell et al. 2015) using BWA-MEM, as implemented in bwa v0.7.10 (Li 2013). Samtools v1.3.1 was used to remove alignments with mapping quality below 20 (Li, et al. 2009). Read-group information including library, lane, sample identity and duplicates was recorded using Picard v2.5 (http://broadinstitute.github.io/picard/). Sequencing reads around insertions and deletions (*i.e*., indels) were realigned using RealignerTargetCreator and IndelRealigner in the Genome Analysis Toolkit (GATK v3.6) (DePristo, et al. 2011). We used the GATK HaplotypeCaller and then GenotypeGVCFs for individual SNP calling and for joint genotyping, respectively, among all samples using default parameters. Finally, we performed several filtering steps using GATK to retain only high-quality SNPs: (1) ‘QD’ < 2.0; (2) ‘FS > 60.0’; (3) ‘MQ < 40.0’; (4) ‘ReadPosRankSum < −8.0’; (5) ‘SOR > 4.0’; (6) ‘MQRankSum < −12.5’.Moreover, we also excluded loci with missing data of more than 30% and discarded two individuals with very low depth of coverage (< 10), as calculated using VCFtools v0.1.15 (http://vcftools.sourceforge.net/man_latest.html). The scripts for SNP calling are available at https://doi.org/10.5281/zenodo.7525386.

### Family relatedness and population structure analysis

To avoid pseudoreplication due to the inclusion of clone mates, we estimated kinship coefficients using the KING toolset for family relationship inference based on pairwise comparisons of SNP data (https://www.kingrelatedness.com/manual.shtml). The software classifies pairwise relationships into four categories according to the estimated kinship coefficient: a negative kinship coefficient estimation indicates the lack of a close relationship. Estimated kinship coefficients higher than >0.354 correspond to duplicates, while coefficients ranging from [0.177, 0.354], [0.0884, 0.177] and [0.0442, 0.0884] correspond to 1^st^-degree, 2^nd^-degree, and 3^rd^-degree relationships, respectively. This analysis identified 13 duplicated genotypes out of a total of 32 samples from the Korean population of *P. davidiana*. In addition, all individuals of *P. qiongdaoensis* were identified as clone mates (Supporting Fig. S23). Therefore, these two populations were eliminated from subsequent analyses and only 7 species were kept for the analyses.

After discarding individuals with low depth and high inbreeding coefficient (F > 0.9, *P. qiongdaoensis*) as well as clones identified with the KING toolset, we used VCFtools v0.1.15 (http://vcftools.sourceforge.net/man_latest.html) to calculate the mean depth of coverage and heterozygosity for each individual. The depth of coverage was relatively homogeneous (Supporting Fig. S24) and varied from 21× to 32×.

We used PLINK (Purcell, et al. 2007) to generate a variance-standardized relationship matrix for principal components analysis (PCA) and a distance matrix to build a neighbor joining tree (NJ-tree) with all filtered SNPs. The NJ tree was constructed using PHYLIP v.3.696 (https://evolution.genetics.washington.edu/phylip.html). Both PCA and NJ-tree analyses were performed based on the full set of SNPs. In addition, we used ADMIXTURE v1.3 for the maximum-likelihood estimation of individual ancestries (Alexander and Lange 2011). First, we generated the input file from a VCF containing unlinked SNPs. Besides, sites with missing data more than 30% have been filtered out. This analysis was run for *K* from 1 to 10, and the estimated parameter standard errors were generated using 200 bootstrap replicates. The best *K* was taken to be the one with the lowest cross-validation error. We also performed an IBD blocks analysis using BEAGLE v5.1 (Browning and Browning 2013) to detect identity-by-descent segments between pairs of species. The parameters we used are: window=100,000; overlap=10,000; ibdtrim=100; ibdlod=10.

### Demographic trajectory reconstruction

To reconstruct the demographic history of *Populus* species, we first inferred the history of species splits and mixture based on genome wide allele frequency data using TreeMix v1.13 (Pickrell and Pritchard 2012). We removed the sites with missing data and performed linkage pruning. We then ran TreeMix implementing a default bootstrap and a block size of 500 SNPs (- k=500). The best migration edge was evaluated according to the greatest increase of total variation explained. The plotting R functions of the TreeMix suite were then used to visualize the results.

### Nucleotide diversity and divergence estimates

Nucleotide diversity, as well as relative and absolute divergence estimates were calculated based on genotype likelihoods. We used ANGSD v0.93 (http://www.popgen.dk/angsd/index.php/ANGSD) to estimate statistical parameters from the BAM files for all *Populus* species. First, we used *‘dosaf 1*’ to calculate site allele frequency likelihood and then used *‘realSFS* to estimate folded site frequency spectra (SFS). Genome-wide diversity and Tajima’s *D* were calculated with the parameter *‘-doThetas 1*’ in ANGSD based on the folded SFS of each species. We selected two population genomic statistics to estimate divergence *F_ST_* and *D_XY_*. We estimated SFS for each population separately and then used it as a prior to generate a 2D-SFS for each species pair. *F_ST_* of each species pair were estimated with the parameters *‘realSFS fst’* based on the 2D-SFS. Finally, we averaged the *F_ST_* value of sites over 10kb windows. To estimate *D_XY_*, we used ANGSD to calculate minor allele frequencies with the parameters *‘-GL 1 -doMaf 1 -only_proper_pairs 1 -uniqueOnly 1 - remove_bads 1 -C 50 -minMapQ 30 -minQ 20 -minInd 4 -SNP_pval 1e-3 -skipTriallelic 1 - doMajorMinor 5* and then computed *D_XY_* as follows: *D_XY_* =A_1_*B_2_+A_2_*B_1_, with A and B being the allele frequencies of A and B, and 1 and 2 being the two populations. We averaged *D_XY_* across 10kb windows.

To examine the relationships among diversity, differentiation, and recombination landscapes, we estimated Pearson’s correlation coefficient between pairs of these statistics. These tests were performed across genomic windows for the 21 possible *Populus* species pairs. Finally, we used *d_a_* (*D_XY_* – mean *π*) as a measure of divergence time.

### Population-scale recombination rate and linkage disequilibrium

We estimated population scaled recombination rate with FastEPRR (Gao, et al. 2016) for each species separately. To eliminate the effect of sample size on the estimation of recombination rate, we downsampled to 13 randomly selected individuals for each species, corresponding to the number of individuals available for *Populus davidiana* (pdav). First, we filtered all missing and non-biallelic sites with VCFtools and then phased the data with the parameters *“impute=true nthreads=20 window=10,000 overlap=1,000 gprobs=false”* in Beagle v5.1 (Browning and Browning 2013). Finally, we ran FastEPRR v2.0 (Gao, et al. 2016) with a window size of 10kb. After getting the results, we estimated the correlation between recombination rate of one species to another. To evaluate LD decay, we used PLINK (Purcell, et al. 2007) to obtain LD statistics for each species. Parameters were set as follows: *‘--maf 0.1 --r^2^ gz --ld-window-kb 500 --ld-window 99999 --ld-window-r^2^ 0*’ LD decay was finally plotted in R.

### Divergent regions of exceptional differentiation

We further investigated genomic differentiation landscapes across multiple species pairs along the *Populus* divergence gradient and identified which evolutionary factors contribute to genomic differentiation. We reported genomic regions showing elevated or decreased values of *F_ST_*, *D_XY_* and *π* across 10kb windows. Windows falling above the top 5% or below the bottom 5% of *F_ST_* and *D_XY_* were considered. For these specific windows, we then classified them following the four models of divergence suggested by Irwin *et al*. 2018 and Han *et al* 2017. These four models differ in the role of gene flow (with or without), or the type of selection (selective sweep, background selection or balancing selection). Note that our detection only considered *F_ST_* and *D_XY_* following the strategy of the two previously mentioned studies, but we encourage investigations with more statistics, including π (as shown in the Box 1) and f_d_ (Supplementary Note 1).

## Acknowledgements

We thank Violaine Llaurens, as well as Camille Roux, Steven van Belleghem and an anonymous reviewer for their careful consideration of this work and their valuable comments on previous versions of this manuscript. We thank members of Jianquan Liu’s lab for collecting samples and to Chris Cole for providing samples from *P. grandidentata*. Michael Barfuss and Elfi Grasserbauer provided assistance in the laboratory. Sequencing was performed at the Next Generation Sequencing Platform of the University of Berne and at the National Genomics Infrastructure (NGI) of Science for Life Laboratory, Stockholm. The Vienna Scientific Cluster (VSC) and the Swedish National Infrastructure for Computing (SNIC) at Uppsala Multidisciplinary Center for Advanced Computational Science (UPPMAX) provided access to computational resources. We are also grateful to members of the PopGen Vienna graduate school for helpful discussions. This manuscript is dedicated to the memory of our friend and colleague Prof. Christian Lexer.

## Funding

This work was supported by a fellowship from the China Scholarship Council (CSC) to Huiying Shang, Swiss National Science Foundation (SNF) grant no.31003A_149306 to Christian Lexer, doctoral programme grant W1225-B20 to a faculty team including Christian Lexer, and the University of Vienna.

## Conflict of interest disclosure

The authors of this preprint declare that they have no conflict of interest with the content of this article.

## Data, script and code availability

The raw read data have been deposited with links to BioProject accession numbers PRJNA299390, PRJNA612655, PRJNA720790, and PRJNA297202 in the NCBI BioProject database (https://www.ncbi.nlm.nih.gov/bioproject/). All the scripts used for the analysis are available on: https://doi.org/10.5281/zenodo.7525386.

## Supplementary information

Supplementary information is available in the “Supplementary material” section of the bioRxiv page of the article, https://doi.org/10.1101/2021.08.26.457771 (latest version).

## Author contributions

Study conceived and designed by Huiying Shang and Christian Lexer. Laboratory work conducted by Huiying Shang. Population genomic data analysis by Huiying Shang with feedback from Thibault Leroy. Interpretation of the results was undertaken by Huiying Shang, Martha Rendón-Anaya, Ovidiu Paun, David Field, Jaqueline Hess, Claus Vogl, Pär K. Ingvarsson, Christian Lexer and Thibault Leroy. The manuscript was drafted by Huiying Shang, with help from Thibault Leroy and Ovidiu Paun, and was improved and approved by all authors.

## Notes

### Competing Interest Statement

The authors have declared no competing interest.

### Summary of Updates

Final version with the PCI Evol Biol badge

