## Supporting information for "Drivers of genomic landscapes of differentiation across *Populus* divergence gradient"

Peer-reviewed & recommended by PCI Evol Biol

**- including Supporting Notes, Tables and Figures -**

[1]Department of Botany and Biodiversity Research, University of Vienna, Vienna, Austria.
[2]Vienna Graduate School of Population Genetics, Vienna, Austria.
[3]Xi'an Botanical Garden, Shaanxi Academy of Sciences, Shaanxi, People's Republic of China.
[4]Swedish University of Agricultural Sciences (SLU), Uppsala, Sweden.
[5]Edith Cowan University, Perth, Australia.
[6]Helmholtz Centre for Environmental Research, Halle (Saale), Germany.
[7]Department of Biomedical Sciences, Vetmeduni Vienna, Vienna, Austria
[8]Key Laboratory for Bio-resources and Eco-environment, College of Life Science, Sichuan University, Chengdu, People's Republic of China
[9]GenPhySE, INRAE, INP, ENVT, Université de Toulouse, Castanet-Tolosan, France
[10]Shared last authorship.
[†]Deceased.

*Corresponding authors:

- Thibault Leroy, GenPhySE, INRAE, INP, ENVT, Université de Toulouse, Castanet-Tolosan, France

- Huiying Shang, Xi'an Botanical Garden, Shaanxi Academy of Sciences, Shaanxi, People's Republic of China.

**Supplementary Note 1：Detection of introgression**

To detect introgression between species at genomic level, we used a modified f-statistics ($f_d$) (Martin et al., 2014) to estimate introgressed sites on non-overlapping 10kb windows with a python script (https://github.com/simonhmartin/genomics_general) among species. The test uses three populations and an outgroup with the relationship (((P1, P2), P3), O) to measure an excess of shared variation between P2 and P3 ($f_d > 0$).

We conducted a correlation analysis between $f_d$ and $F_{ST}$ for the species pairs to test if introgression has a contribution to the accentuated differentiation. The significantly negative correlation between these two parameters supports the role of introgression in shaping the genomic landscape of differentiation, as a result of heterogeneous introgression rates across the genome (Supporting Fig. S6).

**Supplementary Note 2: inferred demographic trajectories of the investigated species**

To infer historical changes in effective population sizes for the investigated *Populus* species we used SMC++ to estimate historical changes in effective population size (Terhorst, et al. 2017), a method which only requires unphased sequence data. To scale coalescent times to real time, a mutation rate of $2.5\times10^{-9}$ per site per year was assumed (Macaya-Sanz, et al. 2012). Due to widespread vegetative reproduction in some poplar species, we have preferred to use a generation time of 20 years rather than the commonly used value of 15 years (Wang, et al. 2016). Given the uncertainties in both the mutation rates and generation times, reported values should not be considered as an exact estimation, but should be rather considered as an order of magnitude.

These inferences (Supporting Fig. S7a) support that all species have experienced population size reductions at least over the last 100,000 years, except for the subtropical species *P. adenopoda* and the North American species *P. tremuloides*. These latter two species have undergone population size expansions from about 30,000 years ago, which seem consistent with their wide present-day distribution areas in South China and North America, respectively (Eckenwalder 1996). Negative mean Tajima's *D* over all non-overlapping 10kb sliding windows spanning the whole genome are consistent with these recent expansions (*P. adenopoda*: -0.97, *P. tremuloides*: -0.61; (Supporting Fig. S7b). For these two species, the folded-SFS also showed a strong excess of rare variants (*P. adenopoda*: 2.4%, *P. tremuloides*: 5.2%; supplementary material, see also Fig. S8), which explains the negative Tajima's *D* values (Supporting Fig. S7b).

### Supplementary Note 3: Selective sweep detection for all populations and species

Selective sweep detection was performed for all species using SweepFinder2 (DeGiorgio *et al.* 2016). To be consistent with all our other summary statistics, SweepFinder2 was run with a 10-kbp space between grid points (-sg 10000). SweepFinder2 was used to calculate a composite likelihood ratio (CLR) to search for signatures of selective sweeps in each species. We reported Manhattan plots of the z-score values for all populations and species (Supporting Fig. S18-S21).

To identify shared or private signatures of sweeps more easily, we then subset regions detected in at least one species and report the number of windows detected in one or more species (Supporting Fig. S22). The vast majority of windows are detected in the single population of each species. This result is consistent with the reduced number of shared regions between species. The comparison exhibiting the greatest number of shared regions is between ptma and ptmae (two populations from the same species *P. tremula*). This species contains at least twice more than any comparison from two (or more) different species. Overall, this result is consistent with the detection of independent selective sweeps and therefore provides additional evidence for the "allopatric selection" scenario.

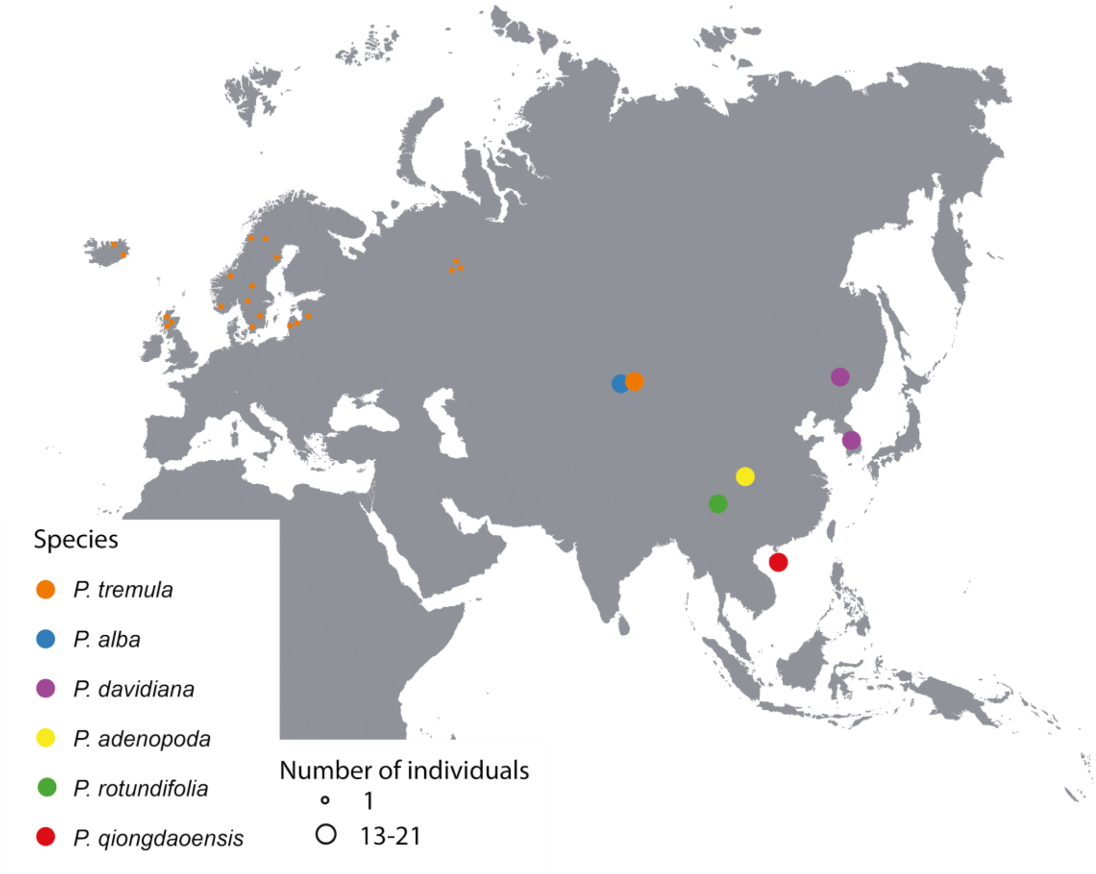


**Supporting Fig. S1.** Map of the *Populus* samples collected in Eurasia. In the map, the different colors represent the different species, while the size of the circle is relative to the amount of samples collected in that location.

*(a)*

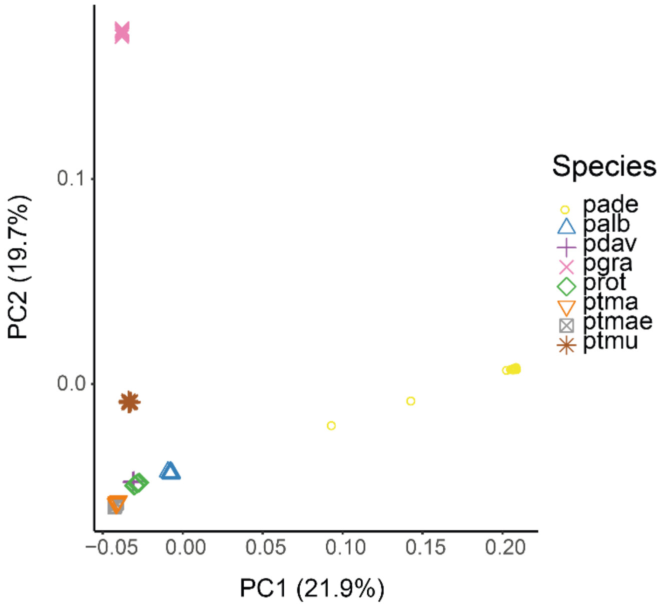


*(b)*

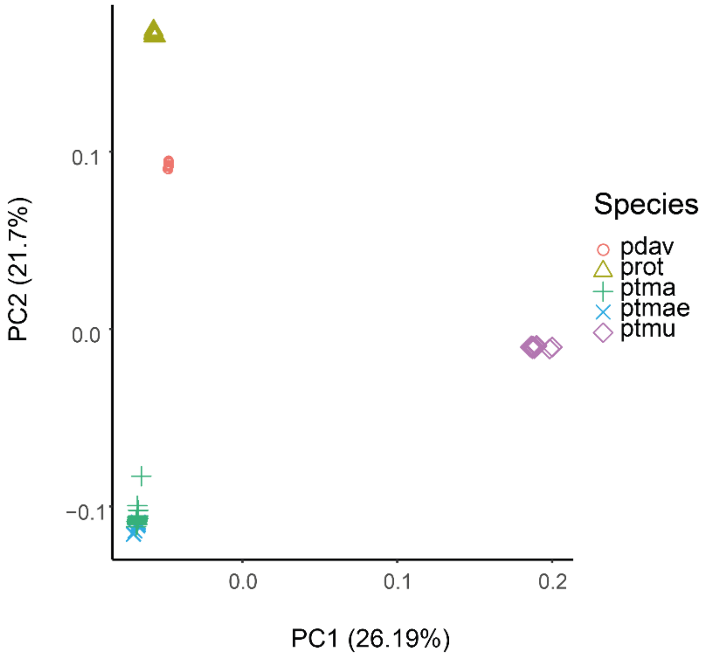


**Supporting Fig. S2.** Two first principal components for a PCA analysis using (a) all resequenced genomes and (b) the four most recently diverged *Populus* species. Species abbreviations: pdav, *P. davidiana* collected in China; prot, *P. rotundifolia*; ptma, *P. tremula* collected in China; ptmae, *P. tremula* collected in Europe; ptmu, *P. tremuloides*.

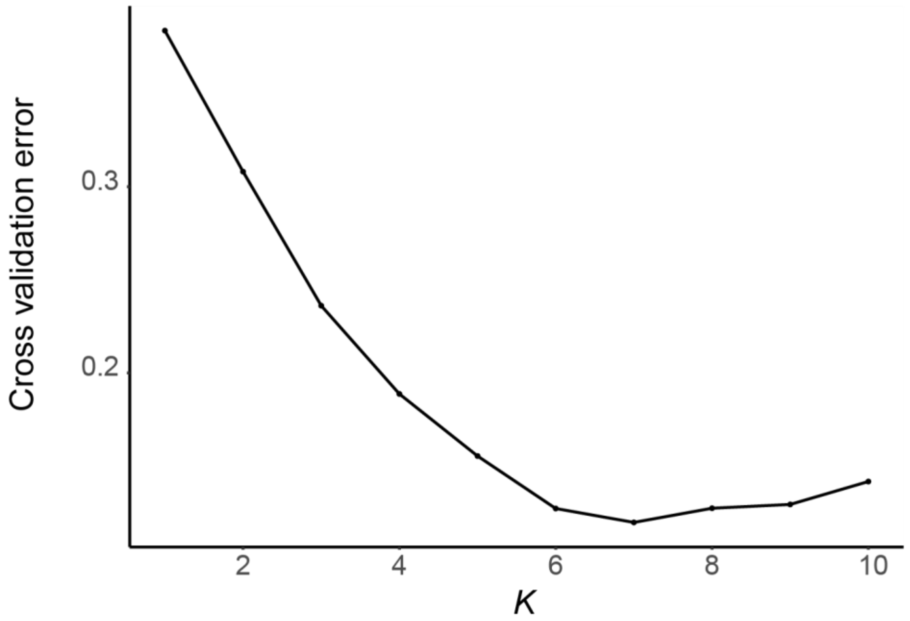


**Supporting Fig. S3.** Plot of ADMIXTURE cross validation error from *K*=1 through *K*=10. The lowest cross validation error obtained at *K*=7.

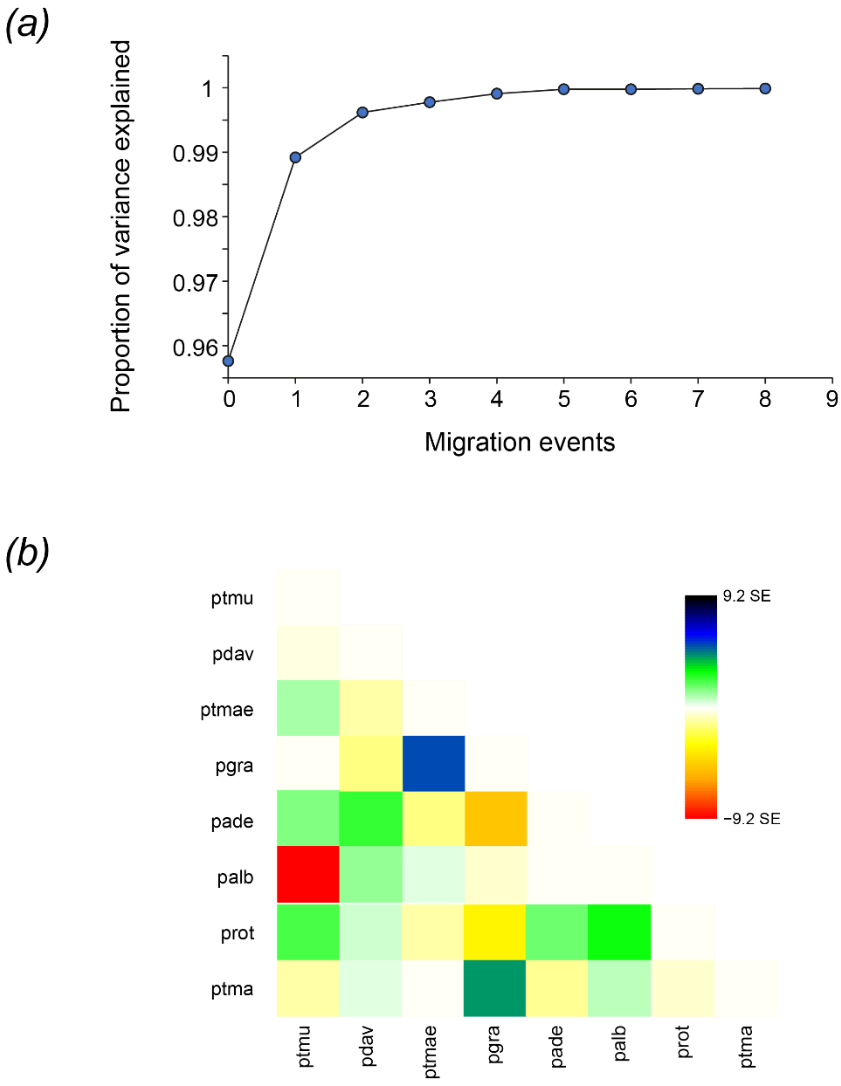


**Supporting Fig. S4.** Treemix analysis results. ***(a)*** plot of proportion of variance explained by the ten models run in TreeMix analysis using m=0 to m=8 migration events. ***(b)*** Residual heatmap from tree with two migration events.

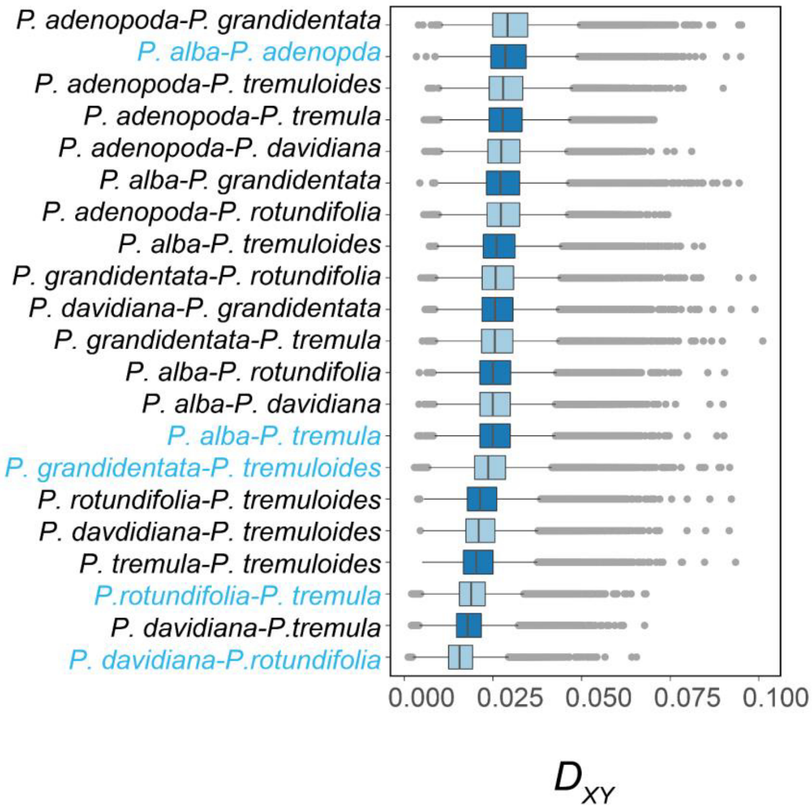


**Supporting Fig. S5.** Observed variance in $D_{XY}$ for all species pairs, *calculated across 10kb windows. The five representative species pairs were labeled in blue.*

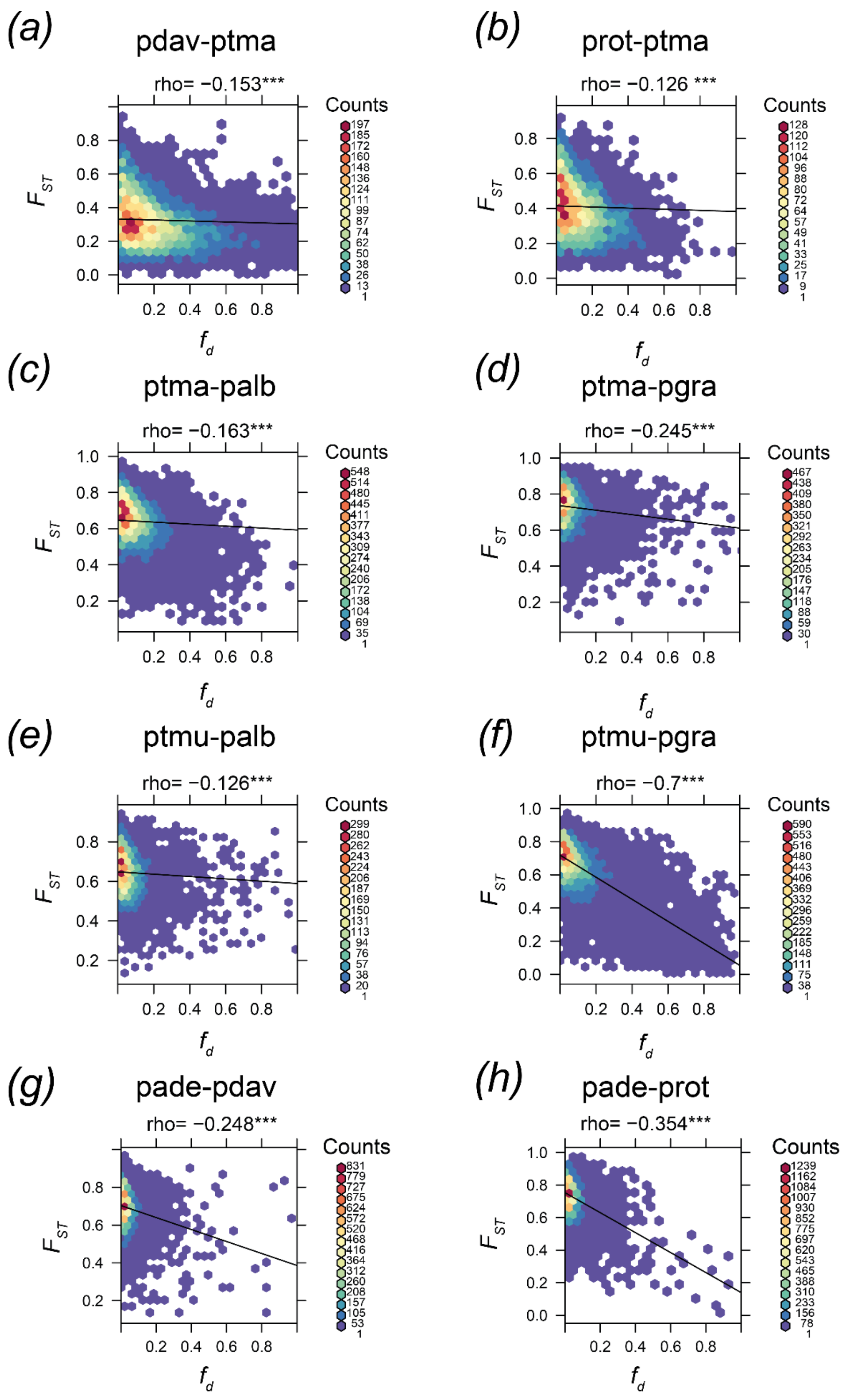


**Supporting Fig. S6.** Correlation analysis between $F_{ST}$ and $f_d$. Species abbreviations: pade, *P. adenopoda*; palb, *P. alba*; pdav, *P. davidiana*; pgra, *P. grandidentata*; prot, *P. rotundifolia*; ptma, *P. tremula*; ptmu, *P. tremuloides*.

(a)

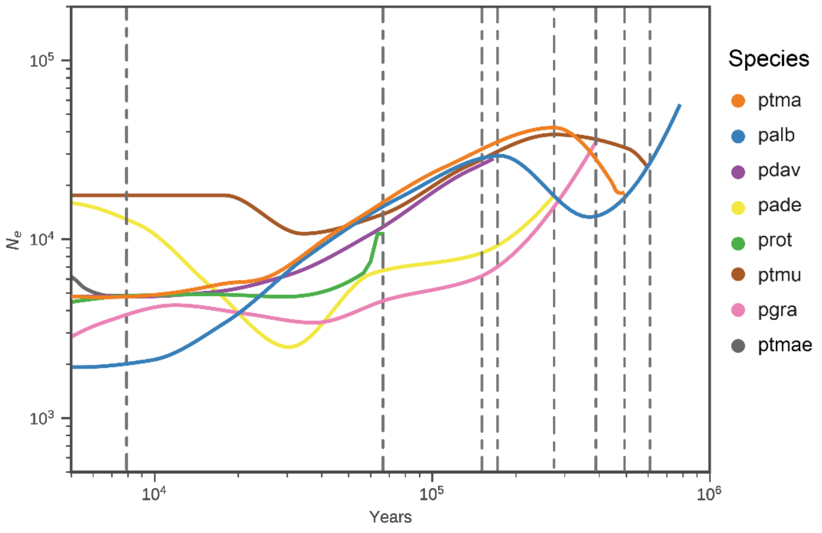


(b)

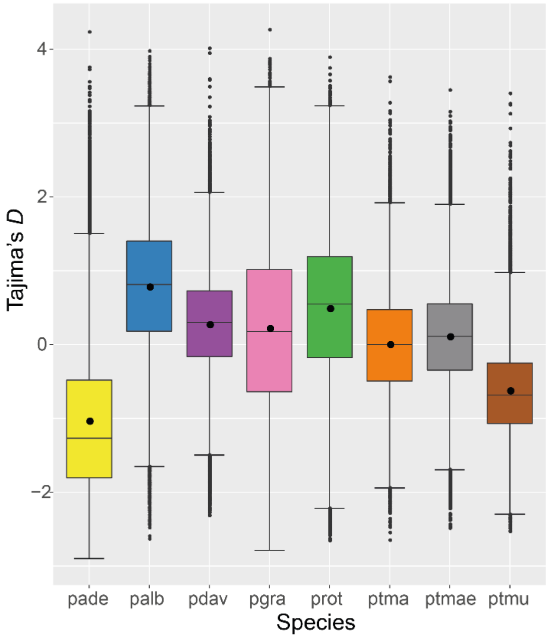


**Supporting Fig. S7.** *Demographic history of Populus species.* ***(a)*** *The maximum likelihood tree inferred by TreeMix under a strictly bifurcating model with two migration events.* ***(b)*** *Changes in effective population size through time inferred by SMC++.* ***(c)*** *The distribution of Tajima's D values calculated over all 10-kb windows.*

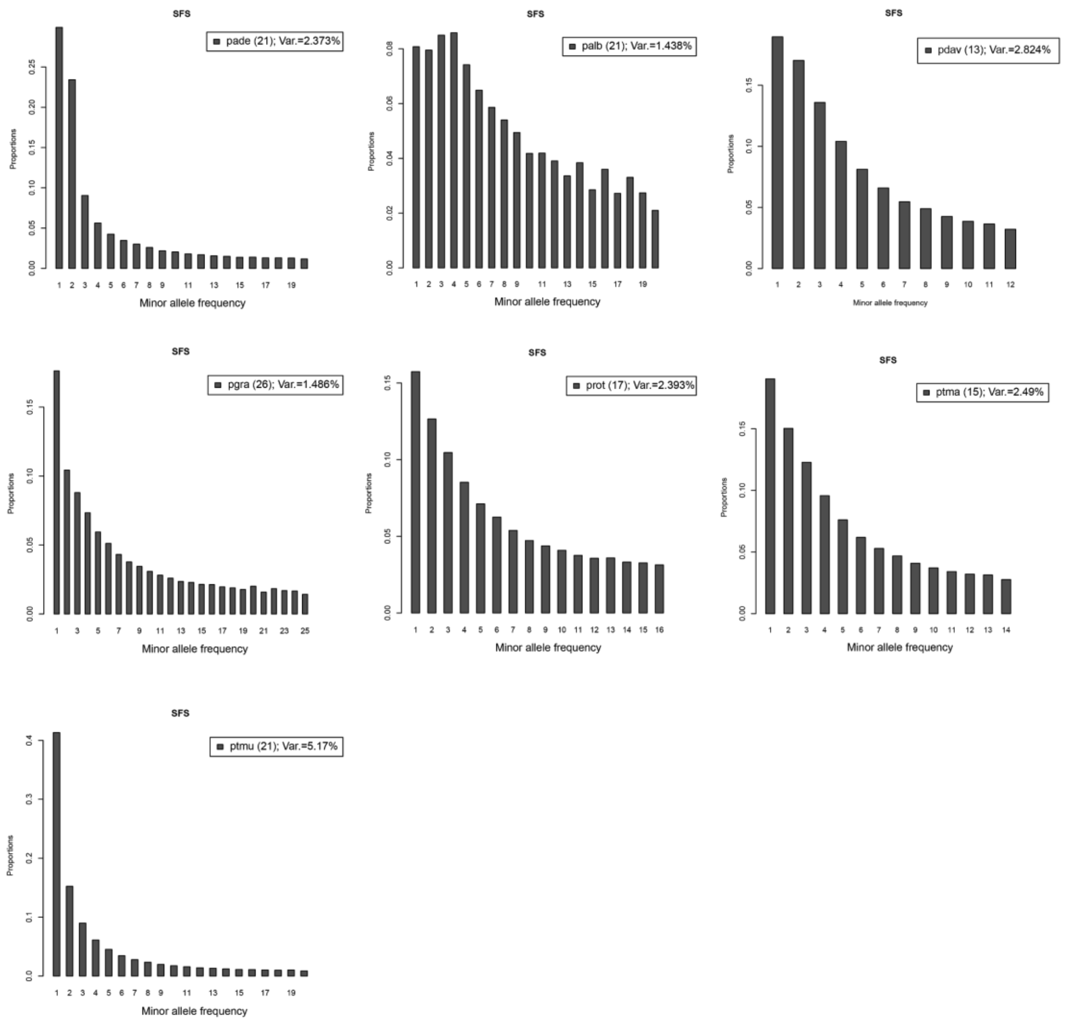


**Supporting Fig. S8.** The folded site frequency spectrum analysis for seven *Populus* species. X-axes show allele accounts in the population, whereas y-axes show the frequency. Species abbreviations: pade, *P. adenopoda*; palb, *P. alba*; pdav, *P. davidiana*; pgra, *P. grandidentata*; prot, *P. rotundifolia*; ptma, *P. tremula*; ptmu, *P. tremuloides*.

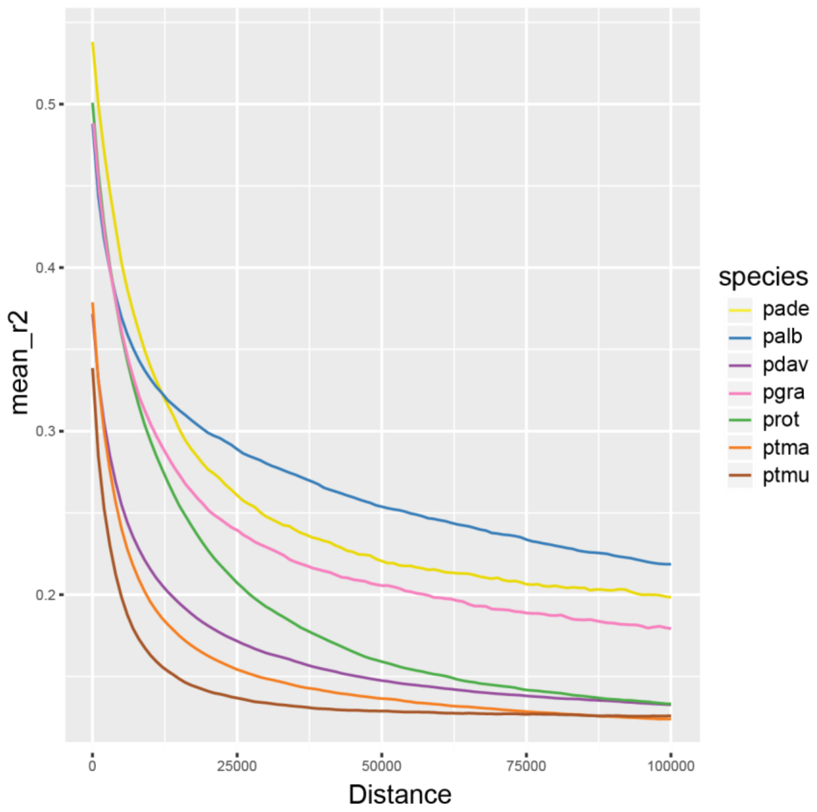


**Supporting Fig. S9.** Linkage disequilibrium (LD) decay by distance across eight *Populus* species. Species abbreviations: pade, *P. adenopoda*; palb, *P. alba*; pdav, *P. davidiana*; pgra, *P. grandidentata*; prot, *P. rotundifolia*; ptma, *P. tremula*; ptmu, *P. tremuloides*

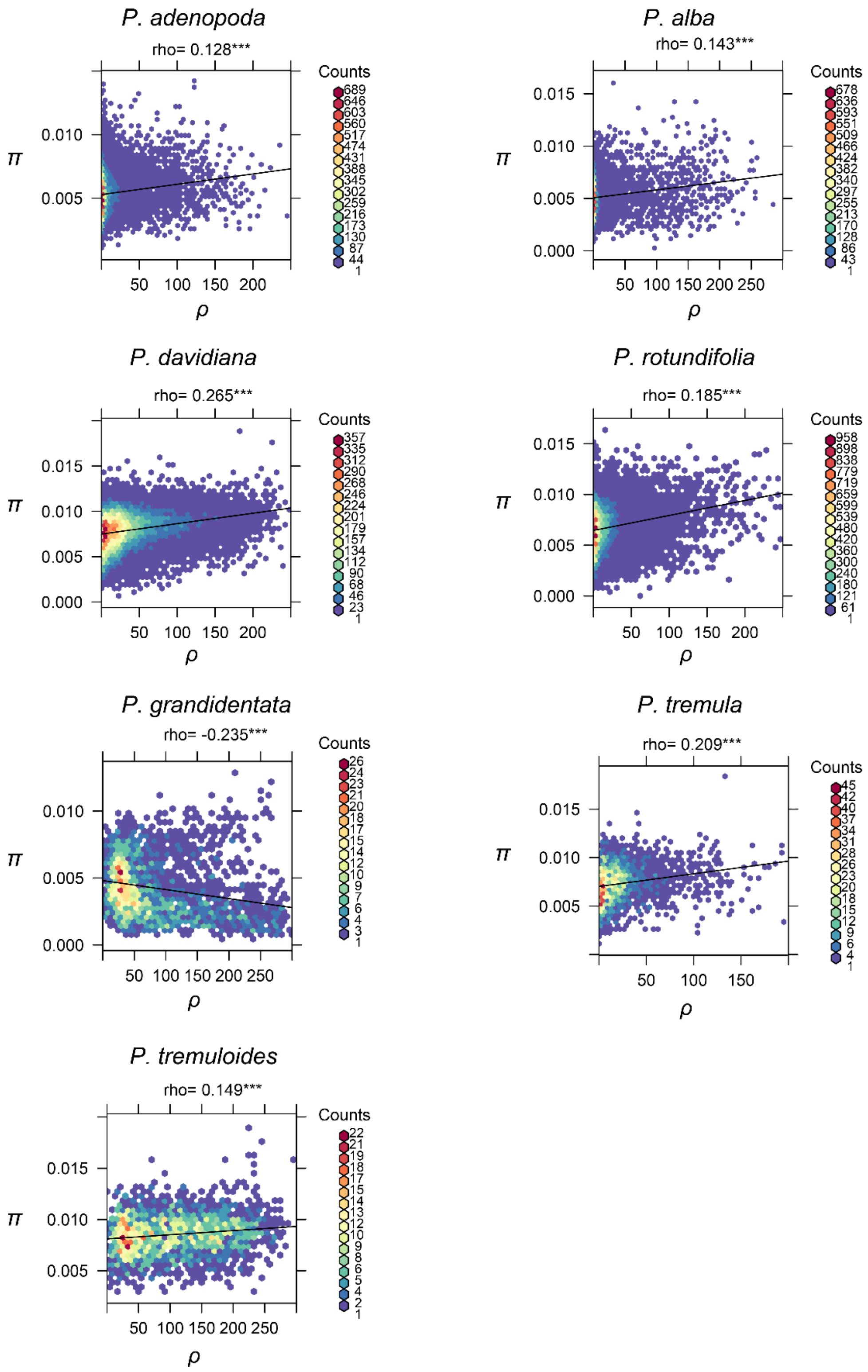


Supporting Fig. S10. Genome-wide correlation analysis between π and ρ for all species. P values less than 0.001 are summarized with three asterisks.

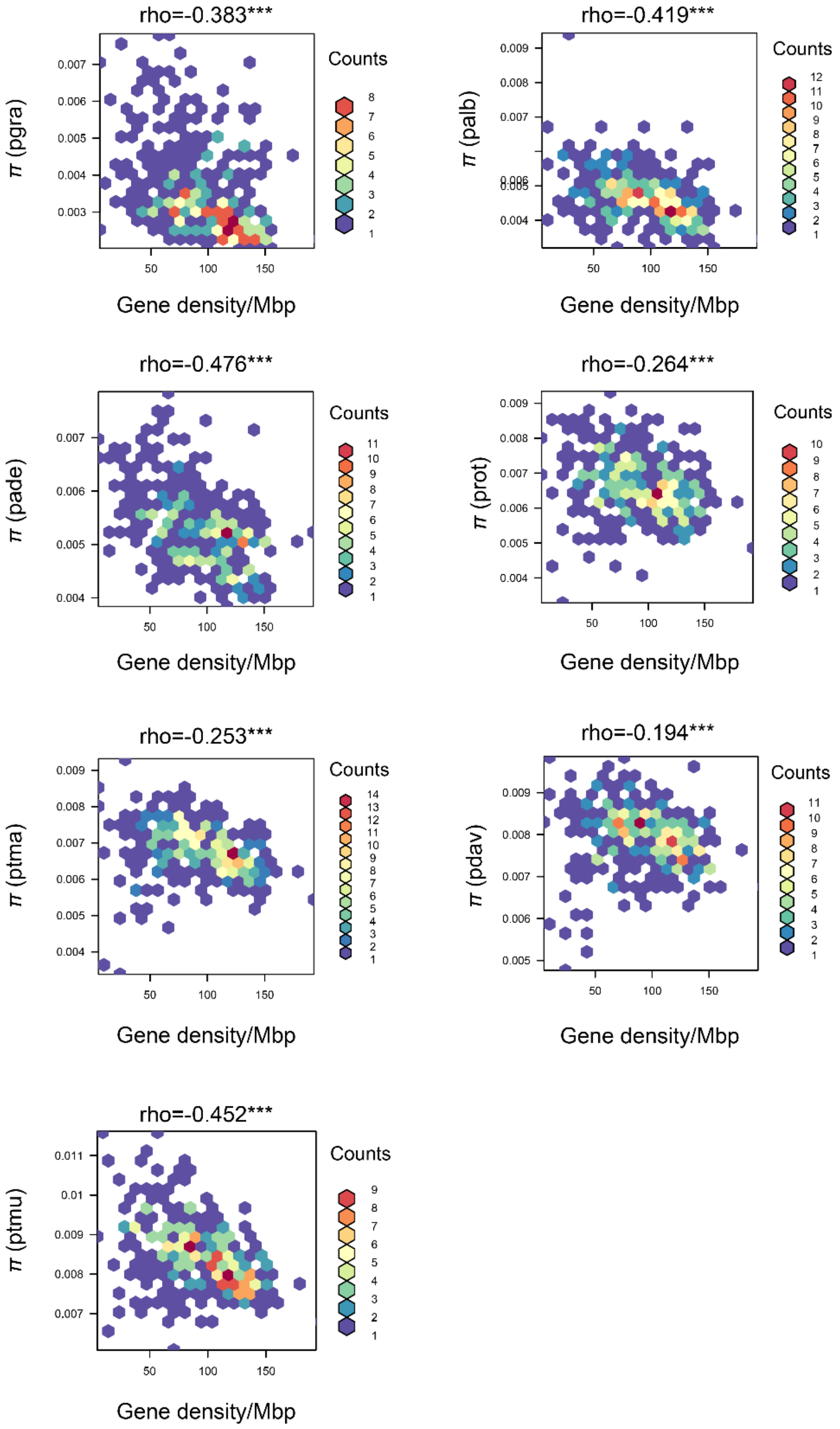


**Supporting Fig. S11.** Correlation analysis between gene density and genetic diversity for each *Populus* species. Species abbreviations: pade, *P. adenopoda*; palb, *P. alba*; pdav, *P. davidiana*; pgra, *P. grandidentata*; prot, *P. rotundifolia*; ptma, *P. tremula*; ptmu, *P. tremuloides*.

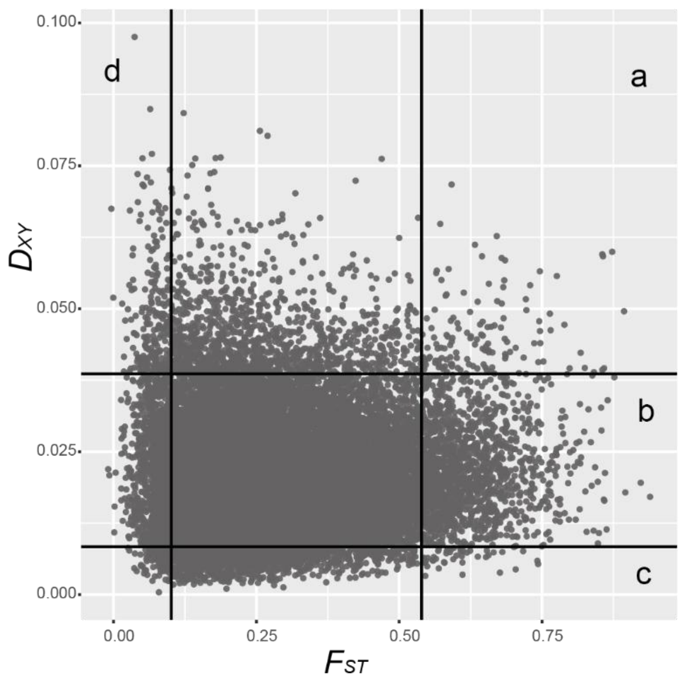


**Supporting Fig. S12.** Speciation models for the formation of genomic islands of relative differentiation. All the grey points were $F_{ST}$ or $D_{XY}$ values based on non-overlapping 10kb windows across the whole genome. The lines were 5% threshold for $F_{ST}$ and $D_{XY}$ values. For example, model (a) explains the regions with top 5% of both $F_{ST}$ and $D_{XY}$. In this model, selection at loci which contribute to reproductive isolation restricts gene exchange between populations, elevating genomic differentiation (lead to higher $F_{ST}$ and $D_{XY}$) and reducing genetic diversity. The second model (b) is 'allopatric selection', selection on distinct regions of the genome after a species split into two populations, leading to lower π and higher $F_{ST}$. As $D_{XY}$ is sensitive to ancestral polymorphism, thus $D_{XY}$ values remain stable in this model. The next model (c) is 'recurrent selection', in this model background selection or selective sweeps at certain regions of the genome reduce genetic diversity in the common ancestor and then selection on the same regions of two daughter populations, leading to lower $D_{XY}$ and $\pi$, higher $F_{ST}$. The last model (d) is 'balancing selection', ancestral polymorphisms are maintained at selected sites, resulting in increased $D_{XY}$ and low $F_{ST}$ between species. The data shown here is for the species pair *P. davidiana*- *P. rotundifolia*, a pair at the early stage of speciation (Fig. 2A).

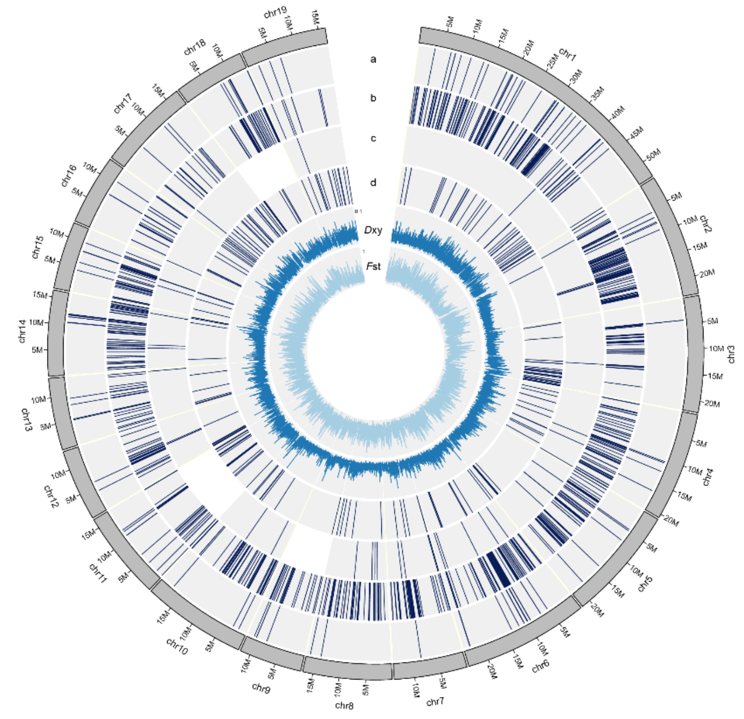


**Supporting Fig. S13.** Genomic regions associated to the four speciation models (a, b, c, and d) across the whole genome in species pair *P. davidiana* – *P. rotundifolia*. $D_{XY}$ and $F_{ST}$ were calculated in 10kb non-overlapping sliding windows.

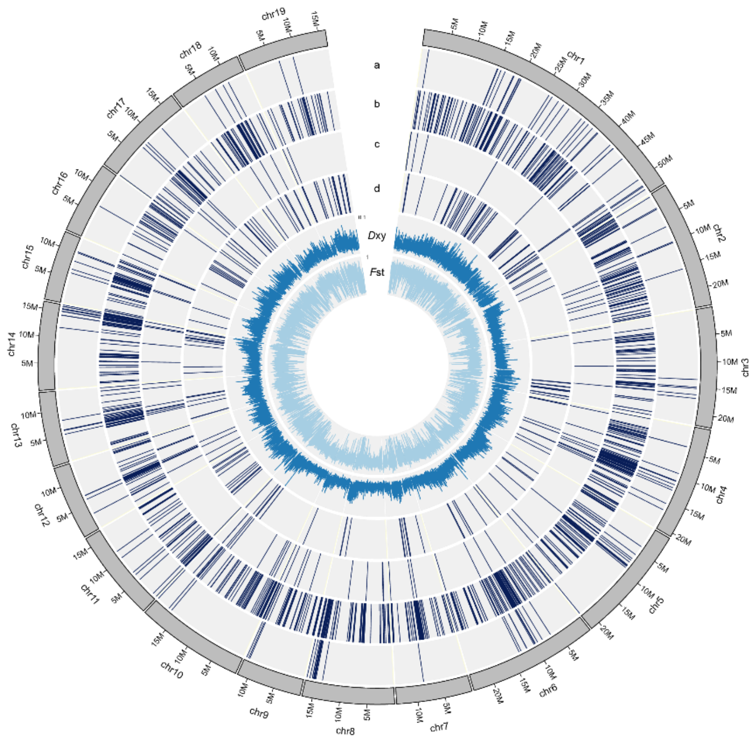


**Supporting Fig. S14.** Genomic regions associated to the four speciation models (a, b, c, and d) across the whole genome in species pair *P. tremula* – *P. alba*. $D_{XY}$ and $F_{ST}$ were calculated in 10kb non-overlapping sliding windows.

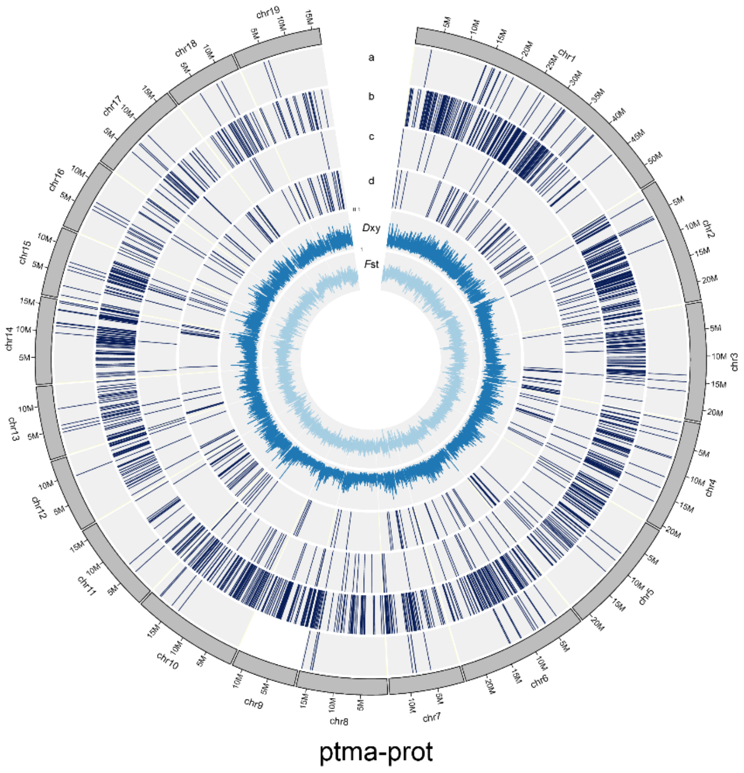


**Supporting Fig. S15.** Genomic regions associated to the four speciation models (a, b, c, and d) across the whole genome in species pair *P. tremula* – *P. rotundifolia*. $D_{XY}$ and $F_{ST}$ were calculated in 10kb non-overlapping sliding windows.

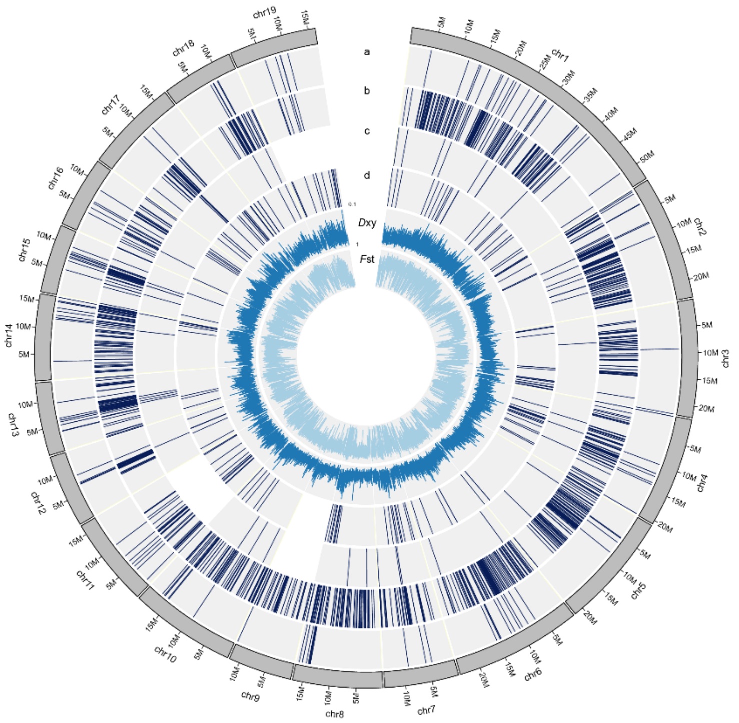


ptmu-pgra

**Supporting Fig. S16.** Genomic regions associated to the four speciation models (a, b, c, and d) across the whole genome in species pair *P. tremuloides – P. grandidentata.* $D_{XY}$ and $F_{ST}$ were calculated in 10kb non-overlapping sliding windows.

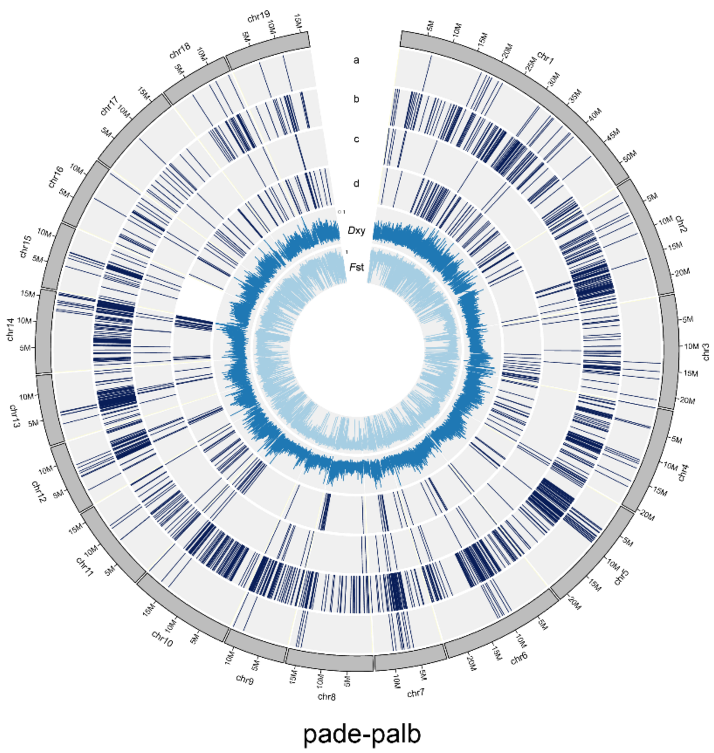


**Supporting Fig. S17.** Genomic regions associated to the four speciation models (a, b, c, and d) across the whole genome in species pair *P. adenopoda* – *P. alba.* $D_{XY}$ and $F_{ST}$ were calculated in 10kb non-overlapping sliding windows.

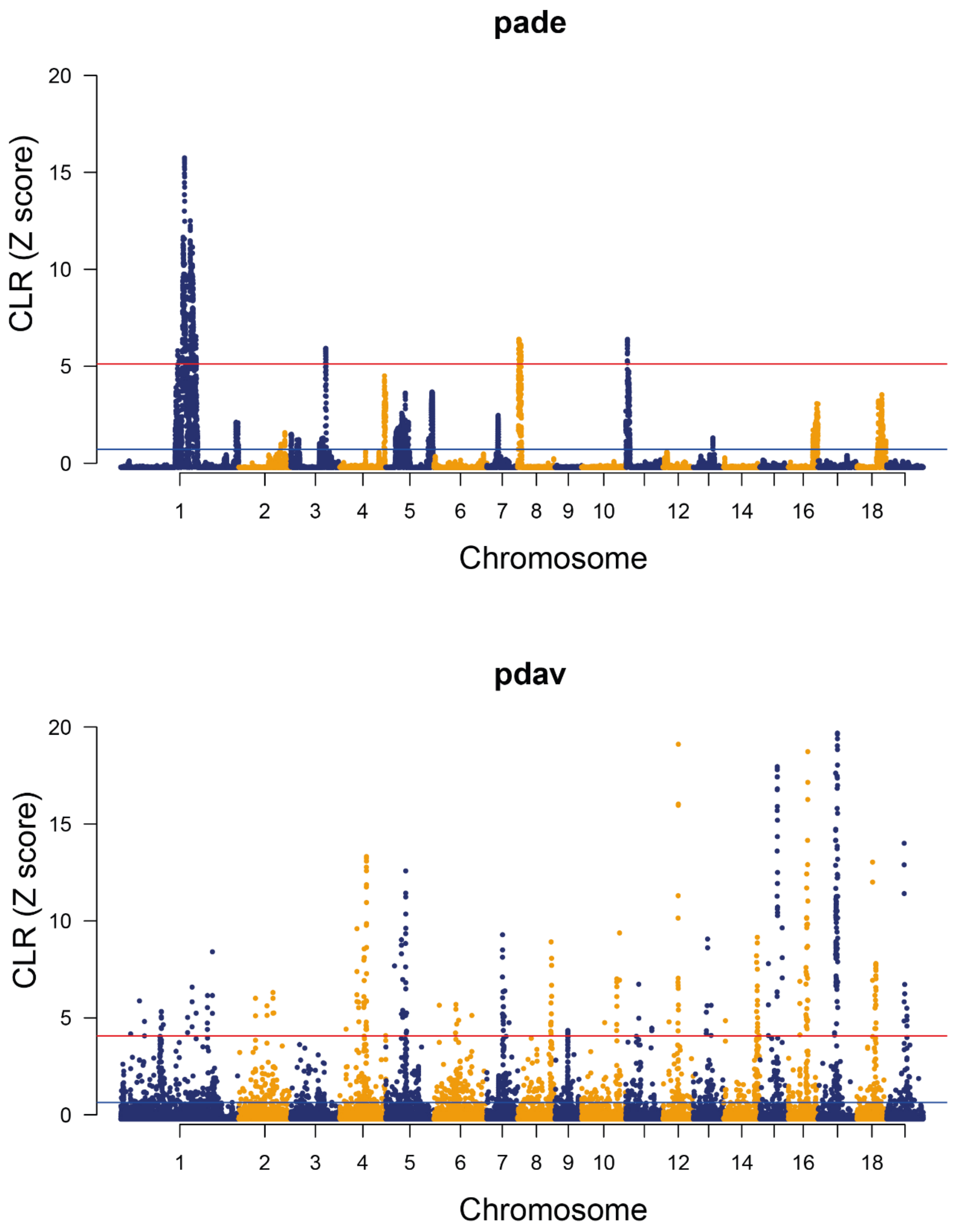


**Supporting Fig. S18.** Genome-wide selective sweeps analysis for SNPs in *P. adenopoda* and *P. davidiana* using composite likelihood ratio (CLR). The red and blue lines are top1% and top5% signatures of selection, respectively.

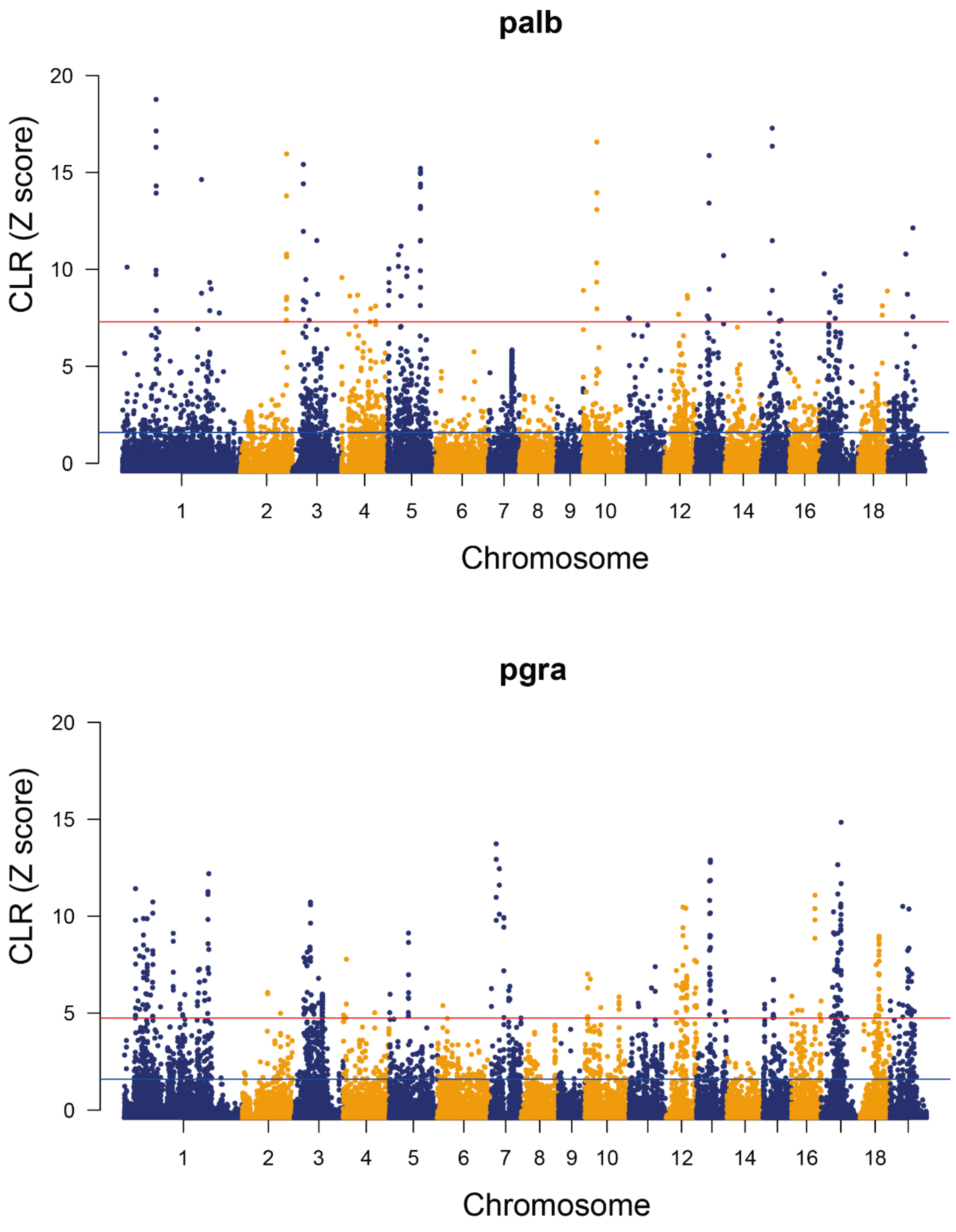


**Supporting Fig. S19.** Genome-wide selective sweeps analysis for SNPs in *P. alba* and *P. grandidentata* using composite likelihood ratio (CLR). The red and blue lines are top1% and top5% signatures of selection, respectively.

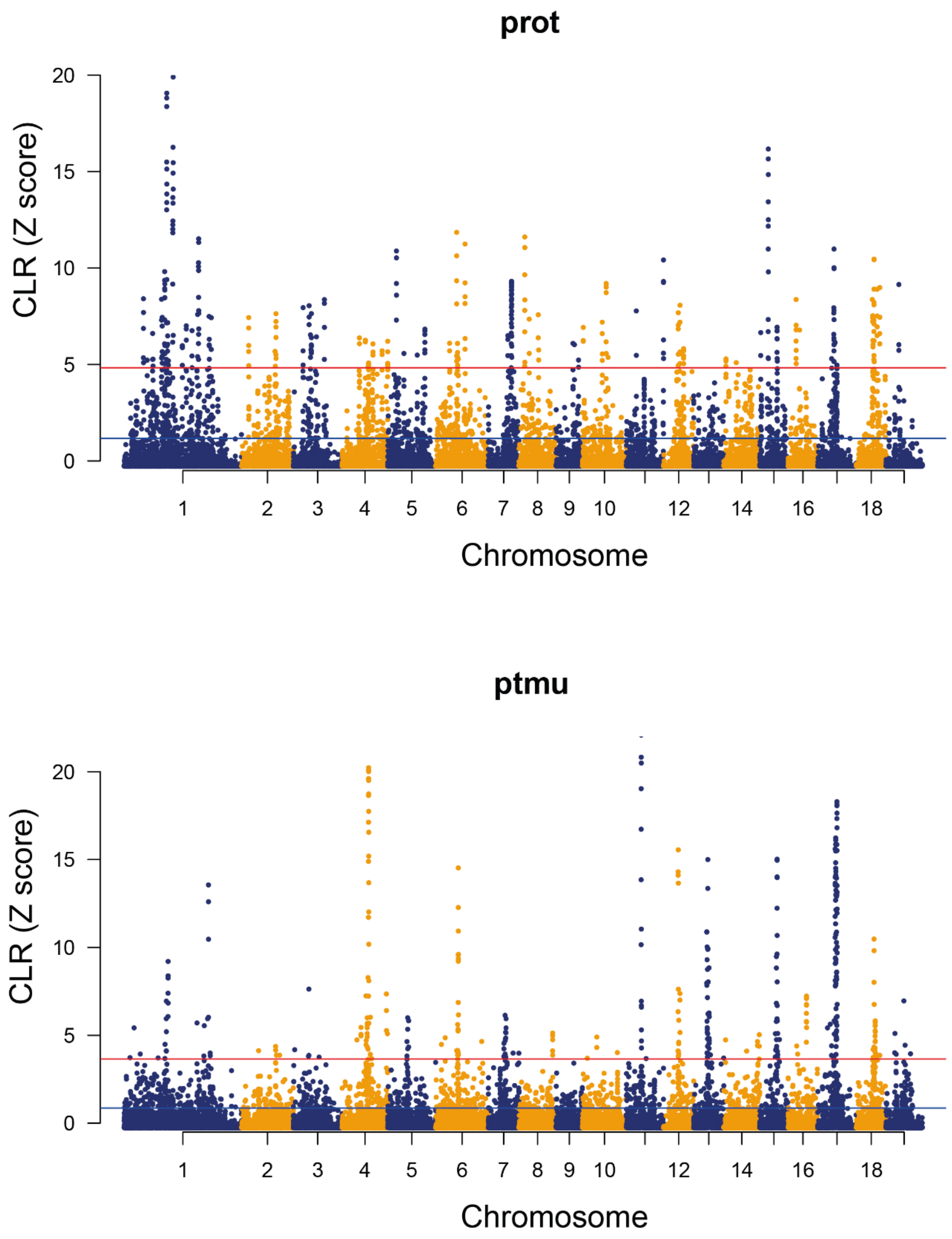


**Supporting Fig. S20.** Genome-wide selective sweeps analysis for SNPs in *P. rotundifolia* and *P. tremuloides* using composite likelihood ratio (CLR). The red and blue lines are top1% and top5% signatures of selection, respectively.

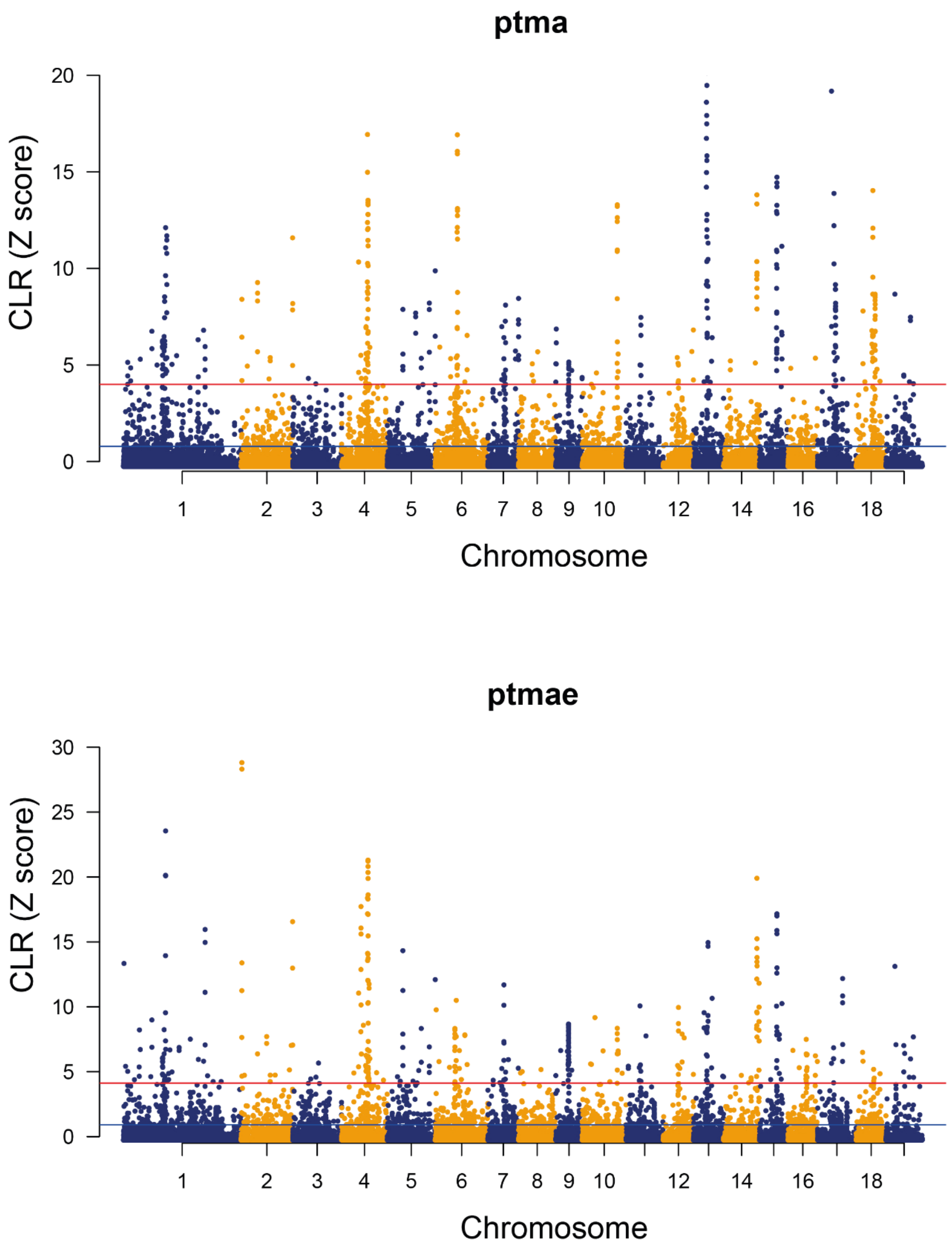


**Supporting Fig. S21.** Genome-wide selective sweeps analysis for SNPs in *P. tremula from China and Europe* using composite likelihood ratio (CLR). The red and blue lines are top1% and top5% signatures of selection, respectively.

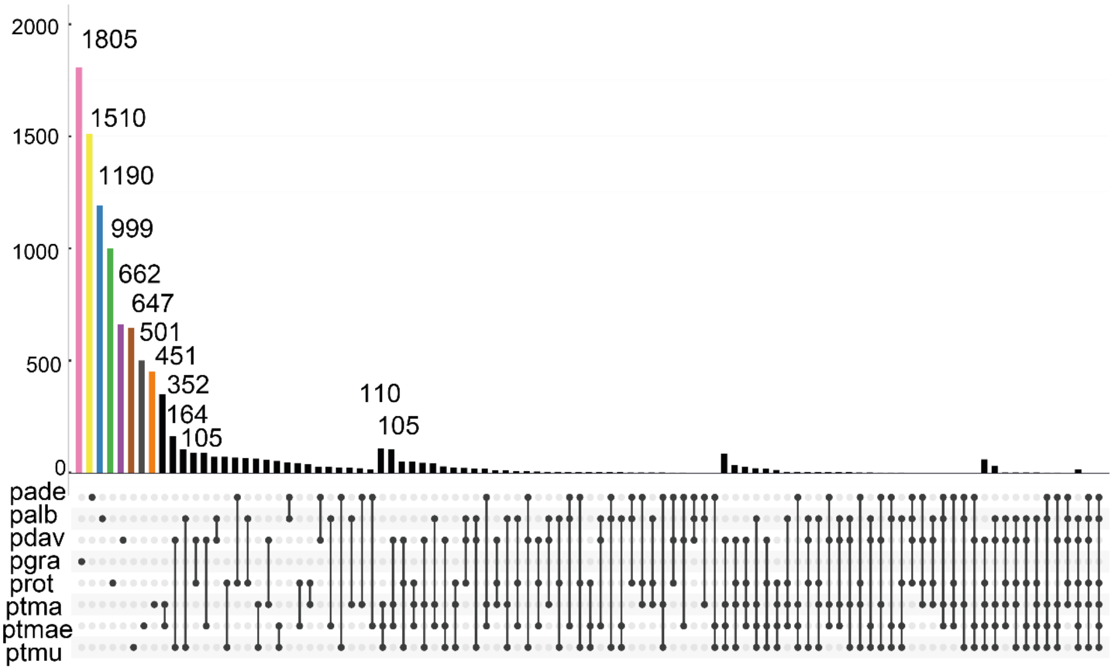


**Supporting Fig. S22.** Number of 10-kbp regions under positive selection for each species. The points and lines show the intersections of the results between species. Above each bar, the number of detected regions is shown, except for those below 100. Species abbreviations: pade, *P. adenopoda*; palb, *P. alba*; pdav, *P. davidiana*; pgra, *P. grandidentata*; prot, *P. rotundifolia*; ptma, *P. tremula* population from China; ptmu, *P. tremuloides*; ptmae, *P. tremula* population from Europe. The plot was drawn using a Python package (https://github.com/hms-dbmi/UpSetR).

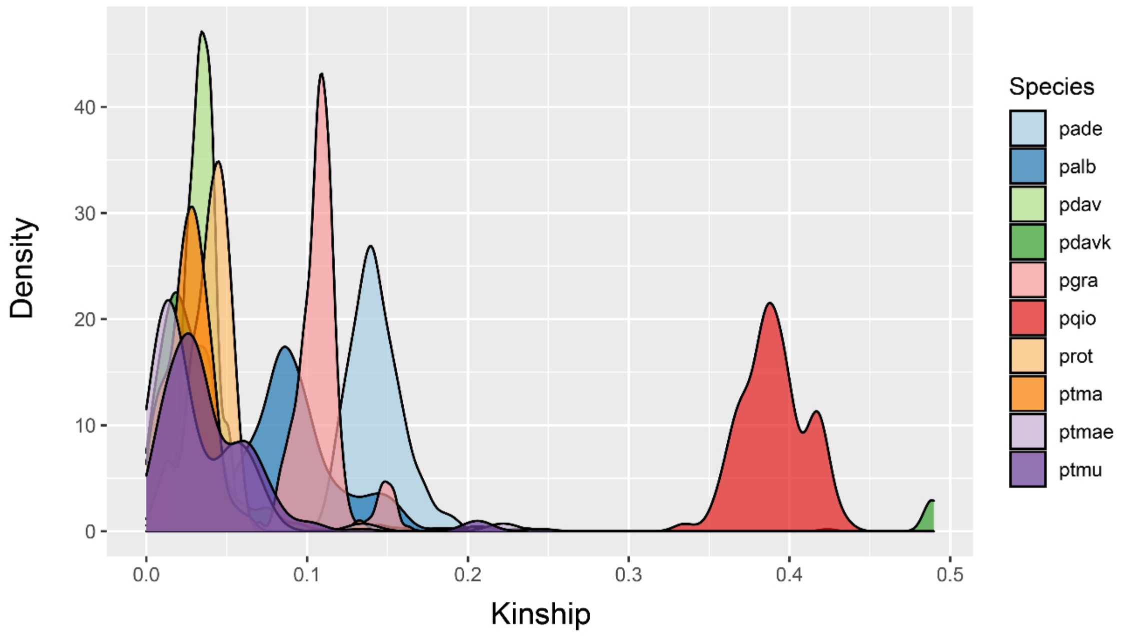


**Supporting Fig. S23.** Family relationship analysis for all eight *Populus* species. Kinship coefficient ranges: >0.354, [0.177, 0.354], [0.0884, 0.177] and [0.0442, 0.0884] corresponding to duplicate/MZ twin, 1st-degree, 2nd-degree, and 3rd-degree relationships, respectively. Species abbreviations: pade, *P. adenopoda*; palb, *P. alba*; pdav, *P. davidiana* collected in China; pdavk, *P. davidiana* collected in South Korea; pgra, *P. grandidentata*; pqio, *P. qiongdaoensis*; prot, *P. rotundifolia*; ptma, *P. tremula* collected in China; ptmae, *P. tremula* collected in Europe; ptmu, *P.tremuloides*.

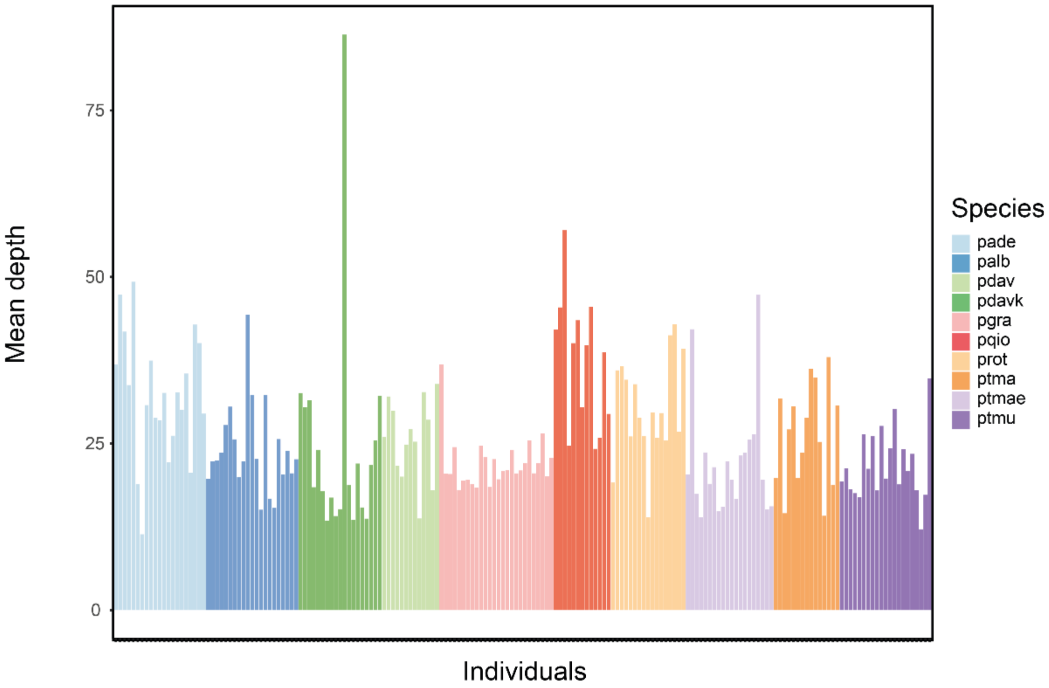


**Supporting Fig. S24.** Mean depth of each *Populus* accession investigated. Species abbreviations: pade, *P. adenopoda*; palb, *P. alba*; pdav, *P. davidiana* collected in China; pdavk, *P. davidiana* collected in South Korea; pgra, *P. grandidentata*; pqio, *P. qiongdaoensis*; prot, *P. rotundifolia*; ptma, *P. tremula* collected in China; ptmae, *P. tremula* collected in Europe; ptmu, *P. tremuloides*.

**Supporting Table S1.** Details of *Populus* samples used in this study.

| Sampled_Trees | Number | Code_number | *Species* | Latitude_(N) | Longitude_(E) | Coverage |
|---|---|---|---|---|---|---|
| pop2014-1 | 21 | palb01 | *Populus alba* | 47.54 | 87.90 | 19.60 |
| pop2014-2 | | palb02 | *Populus alba* | 47.54 | 87.90 | 22.20 |
| pop2014-3 | | palb03 | *Populus alba* | 47.46 | 87.80 | 22.33 |
| pop2014-4 | | palb04 | *Populus alba* | 47.38 | 87.80 | 23.47 |
| pop2014-5 | | palb05 | *Populus alba* | 47.38 | 87.80 | 27.62 |
| pop2014-6 | | palb06 | *Populus alba* | 47.37 | 87.82 | 30.37 |
| pop2014-7 | | palb07 | *Populus alba* | 47.35 | 87.86 | 25.48 |
| pop2014-9 | | palb09 | *Populus alba* | 47.35 | 87.87 | 19.90 |
| pop2014-10 | | palb10 | *Populus alba* | 47.35 | 87.87 | 22.15 |
| pop2014-12 | | palb12 | *Populus alba* | 47.35 | 87.87 | 44.09 |
| pop2014-13 | | palb13 | *Populus alba* | 47.35 | 87.87 | 32.06 |
| pop2014-15 | | palb15 | *Populus alba* | 47.35 | 87.87 | 22.53 |
| pop2014-18 | | palb18 | *Populus alba* | 47.35 | 87.89 | 14.97 |
| pop2014-19 | | palb19 | *Populus alba* | 47.35 | 87.89 | 32.11 |

| | | | | | | |
|---|---|---|---|---|---|---|
| pop2014-20 | | palb20 | *Populus alba* | 47.49 | 87.33 | 16.61 |
| pop2014-21 | | palb21 | *Populus alba* | 47.49 | 86.33 | 15.23 |
| pop2014-24 | | palb24 | *Populus alba* | 47.72 | 86.88 | 25.50 |
| pop2014-26 | | palb26 | *Populus alba* | 47.01 | 86.27 | 20.20 |
| pop2014-37 | | palb37 | *Populus alba* | 47.84 | 86.66 | 23.77 |
| pop2014-38 | | palb38 | *Populus alba* | 47.83 | 86.66 | 20.40 |
| pop2014-41 | | palb41 | *Populus alba* | 47.83 | 86.67 | 22.51 |
| pop2014-45 | 15 | ptma45 | *Populus tremula* | 47.91 | 88.13 | 19.84 |
| pop2014-48 | | ptma48 | *Populus tremula* | 47.96 | 88.18 | 31.73 |
| pop2014-50 | | ptma50 | *Populus tremula* | 47.97 | 88.18 | 14.52 |
| pop2014-52 | | ptma52 | *Populus tremula* | 47.97 | 88.19 | 27.08 |
| pop2014-53 | | ptma53 | *Populus tremula* | 47.98 | 88.20 | 30.50 |
| pop2014-55 | | ptma55 | *Populus tremula* | 47.98 | 88.22 | 19.81 |
| pop2014-56 | | ptma56 | *Populus tremula* | 47.98 | 88.23 | 23.57 |
| pop2014-57 | | ptma57 | *Populus tremula* | 47.98 | 88.24 | 28.84 |
| pop2014-58 | | ptma58 | *Populus tremula* | 47.99 | 88.24 | 36.18 |

| | | | | | | |
|---|---|---|---|---|---|---|
| pop2014-59 | | ptma59 | *Populus tremula* | 47.99 | 88.24 | 34.85 |
| pop2014-62 | | ptma62 | *Populus tremula* | 47.99 | 88.24 | 25.21 |
| pop2014-67 | | ptma67 | *Populus tremula* | 47.99 | 88.24 | 14.13 |
| pop2014-68 | | ptma68 | *Populus tremula* | 48.00 | 88.27 | 37.94 |
| pop2014-70 | | ptma70 | *Populus tremula* | 48.00 | 88.26 | 18.76 |
| pop2014-71 | | ptma71 | *Populus tremula* | 47.99 | 88.26 | 30.67 |
| pop2014-73 | 13 | pdav73 | *Populus davidiana* | 45.32 | 127.35 | 25.89 |
| pop2014-74 | | pdav74 | *Populus davidiana* | 45.10 | 128.00 | 31.94 |
| pop2014-75 | | pdav75 | *Populus davidiana* | 45.02 | 128.18 | 29.83 |
| pop2014-76 | | pdav76 | *Populus davidiana* | 44.95 | 128.79 | 21.57 |
| pop2014-77 | | pdav77 | *Populus davidiana* | 44.93 | 128.97 | 19.95 |
| pop2014-78 | | pdav78 | *Populus davidiana* | 44.92 | 128.99 | 24.74 |
| pop2014-79 | | pdav79 | *Populus davidiana* | 44.53 | 129.79 | 27.09 |
| pop2014-80 | | pdav80 | *Populus davidiana* | 44.77 | 129.15 | 25.18 |
| pop2014-81 | | pdav81 | *Populus davidiana* | 44.93 | 128.73 | 13.71 |
| pop2014-82 | | pdav82 | *Populus davidiana* | 45.25 | 127.73 | 32.63 |

| | | | | | | |
|---|---|---|---|---|---|---|
| pop2014-83 | | pdav83 | *Populus davidiana* | 45.25 | 127.71 | 28.47 |
| pop2014-84 | | pdav84 | *Populus davidiana* | 45.32 | 127.32 | 17.93 |
| pop2014-85 | | pdav85 | *Populus davidiana* | 45.33 | 127.27 | 33.84 |
| LiuJQ-F-2015-01-01 | 21 | pade01 | *Populus adenopoda Maxim.* | 32.76 | 105.25 | 36.63 |
| LiuJQ-F-2015-01-02 | | pade02 | *Populus adenopoda Maxim.* | 32.76 | 105.25 | 47.01 |
| LiuJQ-F-2015-01-03 | | pade03 | *Populus adenopoda Maxim.* | 32.76 | 105.25 | 41.52 |
| LiuJQ-F-2015-01-04 | | pade04 | *Populus adenopoda Maxim.* | 32.76 | 105.25 | 33.50 |
| LiuJQ-F-2015-01-05 | | pade05 | *Populus adenopoda Maxim.* | 32.76 | 105.25 | 48.90 |
| LiuJQ-F-2015-01-06 | | pade06 | *Populus adenopoda Maxim.* | 32.76 | 105.25 | 18.70 |
| LiuJQ-F-2015-01-07 | | pade07 | *Populus adenopoda Maxim.* | 32.76 | 105.25 | 11.26 |
| LiuJQ-F-2015-01-08 | | pade08 | *Populus adenopoda Maxim.* | 32.76 | 105.25 | 30.45 |
| LiuJQ-F-2015-01-09 | | pade09 | *Populus adenopoda Maxim.* | 32.76 | 105.25 | 37.09 |
| LiuJQ-F-2015-01-10 | | pade10 | *Populus adenopoda Maxim.* | 32.76 | 105.25 | 28.67 |
| LiuJQ-F-2015-01-11 | | pade11 | *Populus adenopoda Maxim.* | 32.76 | 105.25 | 28.22 |
| LiuJQ-F-2015-01-12 | | pade12 | *Populus adenopoda Maxim.* | 32.76 | 105.25 | 32.40 |
| LiuJQ-F-2015-01-13 | | pade13 | *Populus adenopoda Maxim.* | 32.76 | 105.25 | 21.95 |
| LiuJQ-F-2015-01-14 | | pade14 | *Populus adenopoda Maxim.* | 32.76 | 105.25 | 25.95 |

| | | | | | | |
|---|---|---|---|---|---|---|
| LiuJQ-F-2015-01-15 | | pade15 | *Populus adenopoda Maxim.* | 32.76 | 105.25 | 32.41 |
| LiuJQ-F-2015-01-16 | | pade16 | *Populus adenopoda Maxim.* | 32.76 | 105.25 | 29.76 |
| LiuJQ-F-2015-01-17 | | pade17 | *Populus adenopoda Maxim.* | 32.76 | 105.25 | 35.21 |
| LiuJQ-F-2015-01-18 | | pade18 | *Populus adenopoda Maxim.* | 32.76 | 105.25 | 20.47 |
| LiuJQ-F-2015-01-19 | | pade19 | *Populus adenopoda Maxim.* | 32.76 | 105.25 | 42.56 |
| LiuJQ-F-2015-01-20 | | pade20 | *Populus adenopoda Maxim.* | 32.76 | 105.25 | 39.80 |
| LiuJQ-F-2015-01-21 | | pade21 | *Populus adenopoda Maxim.* | 32.76 | 105.25 | 29.21 |
| MaoKS-CX-2014-261A-01 | 17 | prot261A01 | *Populus rotundifolia* | 27.14 | 99.39 | 19.06 |
| MaoKS-CX-2014-261A-02 | | prot261A02 | *Populus rotundifolia* | 27.14 | 99.39 | 35.85 |
| MaoKS-CX-2014-261A-03 | | prot261A03 | *Populus rotundifolia* | 27.14 | 99.39 | 36.47 |
| MaoKS-CX-2014-261A-04 | | prot261A04 | *Populus rotundifolia* | 27.14 | 99.39 | 34.46 |
| MaoKS-CX-2014-261A-05 | | prot261A05 | *Populus rotundifolia* | 27.14 | 99.39 | 25.95 |
| MaoKS-CX-2014-261A-06 | | prot261A06 | *Populus rotundifolia* | 27.14 | 99.39 | 33.79 |
| MaoKS-CX-2014-261A-07 | | prot261A07 | *Populus rotundifolia* | 27.14 | 99.39 | 28.73 |
| MaoKS-CX-2014-261A-08 | | prot261A08 | *Populus rotundifolia* | 27.14 | 99.39 | 26.00 |
| MaoKS-CX-2014-261A-09 | | prot261A09 | *Populus rotundifolia* | 27.14 | 99.39 | 13.86 |
| MaoKS-CX-2014-261A-10 | | prot261A10 | *Populus rotundifolia* | 27.14 | 99.39 | 29.59 |

| | | | | | | |
|---|---|---|---|---|---|---|
| MaoKS-CX-2014-261A-11 | | prot261A11 | *Populus rotundifolia* | 27.14 | 99.39 | 25.72 |
| MaoKS-CX-2014-261A-12 | | prot261A12 | *Populus rotundifolia* | 27.14 | 99.39 | 29.44 |
| MaoKS-CX-2014-261A-13 | | prot261A13 | *Populus rotundifolia* | 27.14 | 99.39 | 25.29 |
| MaoKS-CX-2014-261A-14 | | prot261A14 | *Populus rotundifolia* | 27.14 | 99.39 | 41.12 |
| MaoKS-CX-2014-261A-15 | | prot261A15 | *Populus rotundifolia* | 27.14 | 99.39 | 42.80 |
| MaoKS-CX-2014-261A-16 | | prot261A16 | *Populus rotundifolia* | 27.14 | 99.39 | 26.64 |
| MaoKS-CX-2014-261A-17 | | prot261A17 | *Populus rotundifolia* | 27.14 | 99.39 | 39.09 |
| LiuJQ-Tian-2015-001-01 | 14 | pqioT0101 | *Populus qiongdaoensis T.Hong et P.Luo* | 19.12 | 109.09 | 41.81 |
| LiuJQ-Tian-2015-001-02 | | pqioT0102 | *Populus qiongdaoensis T.Hong et P.Luo* | 19.12 | 109.09 | 45.14 |
| LiuJQ-Tian-2015-001-03 | | pqioT0103 | *Populus qiongdaoensis T.Hong et P.Luo* | 19.12 | 109.09 | 56.67 |
| LiuJQ-Tian-2015-001-04 | | pqioT0104 | *Populus qiongdaoensis T.Hong et P.Luo* | 19.12 | 109.09 | 24.46 |
| LiuJQ-Tian-2015-002-01 | | pqioT0201 | *Populus qiongdaoensis T.Hong et P.Luo* | 19.12 | 109.09 | 39.75 |
| LiuJQ-Tian-2015-002-02 | | pqioT0202 | *Populus qiongdaoensis T.Hong et P.Luo* | 19.12 | 109.09 | 43.20 |
| LiuJQ-Tian-2015-002-03 | | pqioT0203 | *Populus qiongdaoensis T.Hong et P.Luo* | 19.12 | 109.09 | 30.17 |
| LiuJQ-Tian-2015-002-04 | | pqioT0204 | *Populus qiongdaoensis T.Hong et P.Luo* | 19.12 | 109.09 | 39.48 |
| LiuJQ-Tian-2015-002-05 | | pqioT0205 | *Populus qiongdaoensis T.Hong et P.Luo* | 19.12 | 109.09 | 45.17 |
| LiuJQ-Tian-2015-002-06 | | pqioT0206 | *Populus qiongdaoensis T.Hong et P.Luo* | 19.12 | 109.09 | 23.98 |

| | | | | | | |
|---|---|---|---|---|---|---|
| LiuJQ-Tian-2015-002-07 | | pqioT0207 | *Populus qiongdaoensis T.Hong et P.Luo* | 19.12 | 109.09 | 25.65 |
| LiuJQ-Tian-2015-002-08 | | pqioT0208 | *Populus qiongdaoensis T.Hong et P.Luo* | 19.12 | 109.09 | 38.38 |
| LiuJQ-Tian-2015-002-09 | | pqioT0209 | *Populus qiongdaoensis T.Hong et P.Luo* | 19.12 | 109.09 | 4.57 |
| LiuJQ-Tian-2015-002-10 | | pqioT0210 | *Populus qiongdaoensis T.Hong et P.Luo* | 19.12 | 109.09 | 29.23 |
| Alb10-3 | 22 | ptmu01 | *P. tremuloides* | NA | NA | 19.18 |
| Alb13-1 | | ptmu02 | *P. tremuloides* | NA | NA | 21.13 |
| Alb16-1 | | ptmu03 | *P. tremuloides* | NA | NA | 18.01 |
| Alb17-4 | | ptmu04 | *P. tremuloides* | NA | NA | 17.45 |
| Alb25-4 | | ptmu05 | *P. tremuloides* | NA | NA | 16.87 |
| Alb27-1 | | ptmu06 | *P. tremuloides* | NA | NA | 26.23 |
| Alb31-1 | | ptmu07 | *P. tremuloides* | NA | NA | 21.09 |
| Alb33-2 | | ptmu08 | *P. tremuloides* | NA | NA | 25.99 |
| Alb35-2 | | ptmu09 | *P. tremuloides* | NA | NA | 17.87 |
| Alb6-3 | | ptmu10 | *P. tremuloides* | NA | NA | 27.43 |
| Albb15-3 | | ptmu11 | *P. tremuloides* | NA | NA | 19.61 |
| Dan1-1C13 | | ptmu12 | *P. tremuloides* | NA | NA | 24.15 |
| Dan2-1B7 | | ptmu13 | *P. tremuloides* | NA | NA | 30.06 |

| | | | | | | |
|---|---|---|---|---|---|---|
| PG1-1B4 | | ptmu14 | *P. tremuloides* | NA | NA | 18.75 |
| PG2-1B9 | | ptmu15 | *P. tremuloides* | NA | NA | 24.03 |
| PG3-1B6 | | ptmu16 | *P. tremuloides* | NA | NA | 20.79 |
| PI12-1B14 | | ptmu17 | *P. tremuloides* | NA | NA | 3.06 |
| PI3-1B3 | | ptmu18 | *P. tremuloides* | NA | NA | 23.28 |
| Sau1-1B10 | | ptmu19 | *P. tremuloides* | NA | NA | 17.88 |
| Sau2-1B2 | | ptmu20 | *P. tremuloides* | NA | NA | 12.05 |
| Sau3-1B13 | | ptmu21 | *P. tremuloides* | NA | NA | 17.20 |
| Wau1-1B5 | | ptmu22 | *P. tremuloides* | NA | NA | 34.64 |
| Bonghyeon4_2 | 32 | pdavk01 | *P. davidiana* | NA | NA | 32.63 |
| Daehwa18-2 | | pdavk02 | *P. davidiana* | NA | NA | 30.40 |
| Daehwa6-1 | | pdavk03 | *P. davidiana* | NA | NA | 30.46 |
| Dongdu2-1 | | pdavk04 | *P. davidiana* | NA | NA | 31.55 |
| KR_BD_5 | | pdavk05 | *P. davidiana* | 37.92 | 128.39 | 18.37 |
| KR_DW_18_2 | | pdavk06 | *P. davidiana* | NA | NA | 23.98 |
| KR_DW_18_3 | | pdavk07 | *P. davidiana* | NA | NA | 17.63 |
| KR_DW_6 | | pdavk08 | *P. davidiana* | 37.50 | 28.46 | 19.13 |

| | | | | | |
|---|---|---|---|---|---|
| KR_DW_6-2 | pdavk09 | *P. davidiana* | 37.50 | 28.46 | 17.77 |
| KR_KM_1 | pdavk10 | *P. davidiana* | 37.29 | 129.24 | 13.41 |
| KR_OD_19-1 | pdavk11 | *P. davidiana* | NA | NA | 16.86 |
| KR_OD_19-2 | pdavk12 | *P. davidiana* | NA | NA | 17.85 |
| KR_OD_19-4 | pdavk13 | *P. davidiana* | 37.80 | 128.54 | 14.05 |
| KR_PD_15 | pdavk14 | *P. davidiana* | 38.16 | 128.37 | 15.10 |
| KR_PG_4 | pdavk15 | *P. davidiana* | 37.83 | 128.49 | 86.26 |
| KR_PG_4-1 | pdavk16 | *P. davidiana* | 37.83 | 128.49 | 14.51 |
| KR_PK_1-2 | pdavk17 | *P. davidiana* | 36.02 | 128.70 | 18.69 |
| KR_PK_2_2 | pdavk18 | *P. davidiana* | NA | NA | 13.50 |
| KR_PK_3_2 | pdavk19 | *P. davidiana* | NA | NA | 21.94 |
| KR_PU-12 | pdavk20 | *P. davidiana* | 37.78 | 128.58 | 15.36 |
| KR_SG_5 | pdavk21 | *P. davidiana* | 37.51 | 128.60 | 24.07 |
| KR_SW_6_2 | pdavk22 | *P. davidiana* | NA | NA | 13.69 |
| KR_SW_6_3 | pdavk23 | *P. davidiana* | NA | NA | 24.99 |
| KR_SY_3 | pdavk24 | *P. davidiana* | 38.18 | 128.30 | 21.73 |

| | | | | | | |
|---|---|---|---|---|---|---|
| KR_SY_4 | | pdavk25 | *P. davidiana* | 38.18 | 128.30 | 25.43 |
| KR_SY_6 | | pdavk26 | *P. davidiana* | 38.18 | 128.30 | 24.02 |
| KR_WD_2_1 | | pdavk27 | *P. davidiana* | NA | NA | 16.55 |
| KR_WD_2_2 | | pdavk28 | *P. davidiana* | NA | NA | 21.26 |
| Palgong1-1 | | pdavk29 | *P. davidiana* | NA | NA | 32.23 |
| Palgong2-3 | | pdavk30 | *P. davidiana* | NA | NA | 27.11 |
| Palgong3-1 | | pdavk31 | *P. davidiana* | NA | NA | 31.31 |
| Sogwang9-1 | | pdavk32 | *P. davidiana* | NA | NA | 37.65 |
| Sgardur | 20 | ptmae01 | *P. tremula* | 65.86 | -17.8929 | 22.28 |
| SJorvik | | ptmae02 | *P. tremula* | 64.84 | -14.37 | 19.57 |
| LV_VIL_02 | | ptmae03 | *P. tremula* | 57.19 | 27.51 | 20.39 |
| LV_SAL_22 | | ptmae04 | *P. tremula* | 56.65 | 22.66 | 42.25 |
| LV_LIE_36 | | ptmae05 | *P. tremula* | 56.24 | 21.40 | 17.42 |
| NO_ALE_07 | | ptmae06 | *P. tremula* | 62.48 | 6.72 | 13.93 |
| NO_STV_02 | | ptmae07 | *P. tremula* | 58.77 | 5.91 | 23.66 |
| NO_MIR_01 | | ptmae08 | *P. tremula* | 66.31 | 14.41 | 18.94 |

| | | | | | | |
|---|---|---|---|---|---|---|
| RU_SYK_01 | | ptmae09 | *P. tremula* | 61.68 | 50.99 | 21.46 |
| RU_SYK_10 | | ptmae10 | *P. tremula* | 61.65 | 51.08 | 14.88 |
| RU_SYK_20 | | ptmae11 | *P. tremula* | 61.67 | 51.09 | 15.48 |
| SwAsp003 | | ptmae12 | *P. tremula* | 56.71 | 13.22 | 16.31 |
| SwAsp033 | | ptmae13 | *P. tremula* | 57.83 | 15.31 | 21.71 |
| SwAsp045 | | ptmae14 | *P. tremula* | 59.64 | 12.94 | 22.08 |
| SwAsp067 | | ptmae15 | *P. tremula* | 61.20 | 13.81 | 23.58 |
| SwAsp096 | | ptmae16 | *P. tremula* | 63.98 | 20.71 | 24.49 |
| SwAsp109 | | ptmae17 | *P. tremula* | 66.36 | 18.18 | 40.91 |
| UK_AAR_23 | | ptmae18 | *P. tremula* | 56.19 | -5.41 | 19.55 |
| UK_CGM_90 | | ptmae19 | *P. tremula* | 57.14 | -3.91 | 15.06 |
| UK_WRS_112 | | ptmae20 | *P. tremula* | 57.87 | -5.44 | 15.55 |
| BT4 | 26 | pgra01 | *P. grandidentata* | NA | NA | 36.91 |
| P11086_106 | | pgra02 | *P. grandidentata* | NA | NA | 20.48 |
| P11086_107 | | pgra03 | *P. grandidentata* | NA | NA | 20.42 |
| P11086_108 | | pgra04 | *P. grandidentata* | NA | NA | 24.47 |

| | | | | | |
|---|---|---|---|---|---|
| P11086_109 | pgra05 | *P. grandidentata* | NA | NA | 18.02 |
| P11086_110 | pgra06 | *P. grandidentata* | NA | NA | 19.40 |
| P11086_111 | pgra07 | *P. grandidentata* | NA | NA | 19.54 |
| P11086_112 | pgra08 | *P. grandidentata* | NA | NA | 18.91 |
| P11086_113 | pgra09 | *P. grandidentata* | NA | NA | 18.39 |
| P11086_114 | pgra10 | *P. grandidentata* | NA | NA | 24.69 |
| P11086_115 | pgra11 | *P. grandidentata* | NA | NA | 22.99 |
| P11086_116 | pgra12 | *P. grandidentata* | NA | NA | 18.48 |
| P11086_117 | pgra13 | *P. grandidentata* | NA | NA | 22.66 |
| P11086_118 | pgra14 | *P. grandidentata* | NA | NA | 19.63 |
| P11086_119 | pgra15 | *P. grandidentata* | NA | NA | 20.87 |
| P11086_120 | pgra16 | *P. grandidentata* | NA | NA | 20.99 |
| P11086_121 | pgra17 | *P. grandidentata* | NA | NA | 24.00 |
| P11086_122 | pgra18 | *P. grandidentata* | NA | NA | 20.52 |
| P11086_123 | pgra19 | *P. grandidentata* | NA | NA | 20.96 |
| P11086_124 | pgra20 | *P. grandidentata* | NA | NA | 22.07 |

| | | | | | |
|---|---|---|---|---|---|
| P11086_125 | pgra21 | *P. grandidentata* | NA | NA | 25.46 |
| P11086_126 | pgra22 | *P. grandidentata* | NA | NA | 20.53 |
| P11086_127 | pgra23 | *P. grandidentata* | NA | NA | 22.01 |
| P11086_128 | pgra24 | *P. grandidentata* | NA | NA | 26.51 |
| P11086_129 | pgra25 | *P. grandidentata* | NA | NA | 20.06 |
| P11086_130 | pgra26 | *P. grandidentata* | NA | NA | 22.88 |

NA, no data.

**Supporting Table S2.** Percentage of each model summarized based on $F_{ST}$ and $D_{XY}$ in each species pair.

| Species_pair | Model | Relative percentage among the detected windows |
|---|---|---|
| pdav-prot | a | 7.2% (129) |
| | b | 78.1% (1396) |
| | c | 2.6% (46) |
| | d | 12.1% (216) |
| ptma-prot | a | 5.5% (98) |
| | b | 75.7% (1357) |
| | c | 7% (125) |
| | d | 11.8% (212) |
| ptma-palb | a | 8.1% (142) |
| | b | 75.7% (1318) |
| | c | 5.2% (92) |
| | d | 11.6% (204) |
| pade-palb | a | 6.9% (121) |
| | b | 74.3% (1313) |
| | c | 4.9% (86) |
| | d | 13.9% (246) |

**Supporting Table S3.** Correlation coefficient between parameters for all species pairs. Species abbreviations: pade, *P. adenopoda*; palb, *P. alba*; pdav, *P. davidiana*; pgra, *P. grandidentata*; prot, *P. rotundifolia*; ptma, *P. tremula* population from China; ptmu, *P. tremuloides*; ptmae, *P. tremula* population from Europe.

| Species pairs | $d_a$ | $\pi$ & $F_{ST}$ | $\pi$ & $D_{XY}$ | $\pi$ & $\rho$ | $F_{ST}$ & $\rho$ | $F_{ST}$ & $D_{XY}$ | $D_{XY}$ & $\rho$ |
|---|---|---|---|---|---|---|---|
| pdav-prot | 0.009108 | -0.373 | 0.502 | 0.28 | -0.243 | 0.123 | 0.145 |
| pdav-ptma | 0.01138 | -0.428 | 0.408 | 0.199 | -0.247 | 0.222 | 0.0463 |
| prot-ptma | 0.013101 | -0.53 | 0.333 | 0.195 | -0.204 | 0.206 | 0.147 |
| ptma-ptmu | 0.01404 | -0.566 | 0.283 | 0.148 | -0.149 | 0.332 | 0.0936 |
| pdav-ptmu | 0.014144 | -0.558 | 0.31 | 0.272 | -0.245 | 0.314 | 0.0609 |
| prot-ptmu | 0.015303 | -0.63 | 0.272 | 0.113 | -0.0795 | 0.261 | 0.147 |
| pgra-ptmu | 0.019167 | -0.71 | 0.123 | 0.0432 | 0.113 | 0.292 | 0.0212 |
| palb-pdav | 0.02014 | -0.616 | 0.231 | 0.251 | -0.3 | 0.189 | 0.207 |
| palb-ptma | 0.020623 | -0.586 | 0.239 | 0.212 | -0.252 | 0.202 | 0.197 |
| palb-prot | 0.020878 | -0.686 | 0.186 | 0.207 | -0.314 | 0.143 | 0.205 |
| palb-ptmu | 0.021171 | -0.7 | 0.256 | 0.0997 | -0.0502 | 0.154 | 0.202 |
| pdav-pgra | 0.021413 | -0.709 | 0.131 | 0.349 | 0.214 | 0.171 | 0.318 |
| pgra-ptma | 0.021838 | -0.694 | 0.147 | 0.201 | -0.219 | 0.173 | 0.317 |

| | | | | | | | |
|---|---|---|---|---|---|---|---|
| pgra-prot | 0.02224 | -0.768 | 0.0851 | 0.249 | -0.33 | 0.143 | 0.322 |
| pade-pdav | 0.022243 | -0.535 | 0.265 | 0.261 | -0.326 | 0.202 | 0.222 |
| pade-ptmu | 0.022809 | -0.645 | 0.292 | 0.0673 | -0.00495 | 0.155 | 0.22 |
| pade-prot | 0.022838 | -0.613 | 0.206 | -0.00308 | 0.0786 | 0.184 | 0.224 |
| pade-ptma | 0.023245 | -0.509 | 0.289 | 0.185 | -0.234 | 0.199 | 0.217 |
| palb-pgra | 0.024776 | -0.773 | 0.167 | 0.059 | -0.207 | 0.0425 | 0.178 |
| pade-palb | 0.025411 | -0.628 | 0.279 | 0.0768 | -0.244 | 0.0949 | 0.213 |
| pade-pgra | 0.026536 | -0.742 | 0.174 | 0.111823 | -0.0876 | 0.0853 | 0.204 |

**Supporting Table S4.** Mapping region to the reference genome.

| Species | Mapping regions length | Genome coverage |
|---|---|---|
| Reference genome(ptmae) | 381973357 | - |
| palb | 306982824 | 80% |
| pade | 305905287 | 80% |
| pdav | 334266980 | 88% |
| prot | 324497334 | 85% |
| pgra | 322119691 | 84% |
| pdavk | 326008035 | 85% |
| ptma | 351988399 | 92% |
| ptmae | 352182536 | 92% |
| ptmu | 330819811 | 87% |
| pqio | 305125633 | 80% |
